# Hippocampal and cortical mechanisms at retrieval explain variability in episodic remembering in older adults

**DOI:** 10.1101/2020.02.11.940668

**Authors:** Alexandra N. Trelle, Valerie A. Carr, Scott A. Guerin, Monica K. Thieu, Manasi Jayakumar, Wanjia Guo, Ayesha Nadiadwala, Nicole K. Corso, Madison P. Hunt, Celia P. Litovsky, Natalie J. Tanner, Gayle K. Deutsch, Jeffrey D. Bernstein, Marc B. Harrison, Anna M. Khazenzon, Jiefeng Jiang, Sharon J. Sha, Carolyn A. Fredericks, Brian K. Rutt, Elizabeth C. Mormino, Geoffrey A. Kerchner, Anthony D. Wagner

## Abstract

Age-related episodic memory decline is characterized by striking heterogeneity across individuals. Hippocampal pattern completion is a fundamental process supporting episodic memory. Yet, the degree to which this mechanism is impaired with age, and contributes to variability in episodic memory, remains unclear. We combine univariate and multivariate analyses of fMRI data from a large cohort of cognitively normal older adults (N=100; 60-82 yrs) to measure hippocampal activity and cortical reinstatement during retrieval of trial-unique associations. Trial-wise analyses revealed that hippocampal activity predicted cortical reinstatement strength, and these two metrics of pattern completion independently predicted retrieval success. However, increased age weakened cortical reinstatement and its relationship to memory behaviour. Critically, individual differences in the strength of hippocampal activity and cortical reinstatement explained unique variance in performance across multiple assays of episodic memory. These results indicate that fMRI indices of hippocampal pattern completion explain within- and across-individual memory variability in older adults.

---

Episodic memory – in particular the ability to form and retrieve associations between multiple event elements that comprise past experiences – declines with age (1–3). Retrieval of an episodic memory relies critically on hippocampal-dependent pattern completion, which entails reactivation of a stored memory trace by the hippocampus in response to a partial cue, leading to replay of cortical activity patterns that were present at the time of memory encoding (4–7). Given observed links between in vivo measures of pattern completion and episodic remembering (8–10), and evidence of altered hippocampal function with age (11–12), changes in hippocampal pattern completion may play an important role in explaining age-related impairments in episodic memory. While a leading hypothesis, the degree to which the integrity of pattern completion can explain (a) trial-to-trial differences in episodic remembering within older adults and (b) differences in memory performance between older individuals remain underspecified.

Functional MRI (fMRI) studies in younger adults suggest that hippocampal pattern completion is associated with at least two key neural markers: (a) an increase in hippocampal univariate activity (13–15) and (b) cortical reinstatement of content-specific activity patterns present during encoding (16–18). Multivariate pattern analyses –– machine learning classification (19) and pattern similarity (20) –– reveal evidence for cortical reinstatement of categorical event features (10, 21–22) and event-specific details (23–25) during successful recollection. Moreover, hippocampal and cortical metrics of pattern completion covary, such that trial-wise fluctuations in hippocampal univariate retrieval activity predict the strength of cortical reinstatement (10, 23–24), and both hippocampal activity and reinstatement strength predict associative retrieval performance (10, 26). These findings support models (4–6) positing that cortical reinstatement depends, in part, on hippocampal processes, and contributes to remembering.

Initial data bearing on age-related changes in hippocampal pattern completion are mixed. Studies comparing hippocampal activity during episodic retrieval in older and younger adults have revealed age-related reductions in activity (27, 28) and age-invariant effects (29, 30). Similarly, while some have identified reduced category-level (31, 32) and event-level (33, 34) cortical reinstatement in older relative to younger adults, others observed age-invariant category-level reinstatement (29) or that age-related differences in reinstatement strength are eliminated after accounting for the strength of category representations during encoding (35). Although extant studies have yielded important initial insights, the absence of trial-wise analyses relating hippocampal activity to cortical reinstatement, or relating each of these neural measures to memory behaviour, prevents clear conclusions regarding the degree to which hippocampal pattern completion processes are impacted with age. Aging may affect one or both of these neural processes, and/or may disrupt the predicted relationships between these neural variables and behaviour (e.g., 10). The first aim of the present study is to quantify trial-wise fluctuations in hippocampal activity and cortical reinstatement in older adults, and examine how these measures relate to one another, as well as how these measures relate to episodic remembering of trial-unique associative content.

Critically, in addition to varying within individuals, the degree to which pattern completion processes are disrupted among older adults may vary across individuals. Indeed, age-related memory decline is characterized by striking heterogeneity, with some individuals performing as well as younger adults and others demonstrating marked impairment (36-37, see 38 for review). Identifying the neural factors driving this variability is a clear emerging aim of cognitive aging research (38, 40). However, due to modest sample sizes, extant studies typically lack sufficient power to examine individual differences in retrieval mechanisms among older adults (28–35). Moreover, while recent work examining variability in hippocampal function has demonstrated relationships between hippocampal retrieval activity and associative memory performance in older adults (36, 39), the direction of this relationship differed across studies; to date, the relationship between individual differences in cortical reinstatement and memory performance remains unexplored. As such, the second aim of the present study is to examine whether hippocampal and cortical indices of pattern completion vary with age, and to assess the degree to which these measures explain individual differences in episodic memory performance –– both as a function of age and independent of age.

To address these two aims, a large sample (N=100) of cognitively normal older participants (60-82 yrs) from the Stanford Aging and Memory Study (SAMS; **Table 1; Methods**) performed an associative memory task (**Figure 1**) concurrent with high-resolution fMRI. Participants intentionally studied trial-unique word-picture pairs (concrete nouns paired with famous faces and famous places), and then had their memory for the word-picture associations probed. During retrieval scans, participants viewed a studied or novel word on each trial and indicated whether they (a) recollected the associate paired with the word, responding ‘face’ or ‘place’ accordingly (providing an index of associative memory), (b) recognized the word as ‘old’ but were unable to recall the associate (providing an index of item memory –– putatively reflecting familiarity, non-criterial recollection, or a mix of the two), or (c) thought the word was ‘new’. Following scanning, participants were shown the studied words again and asked to recall the specific associate paired with each word, this time explicitly providing details of the specific image (providing an index of exemplar-specific recall).

**Figure 1.**
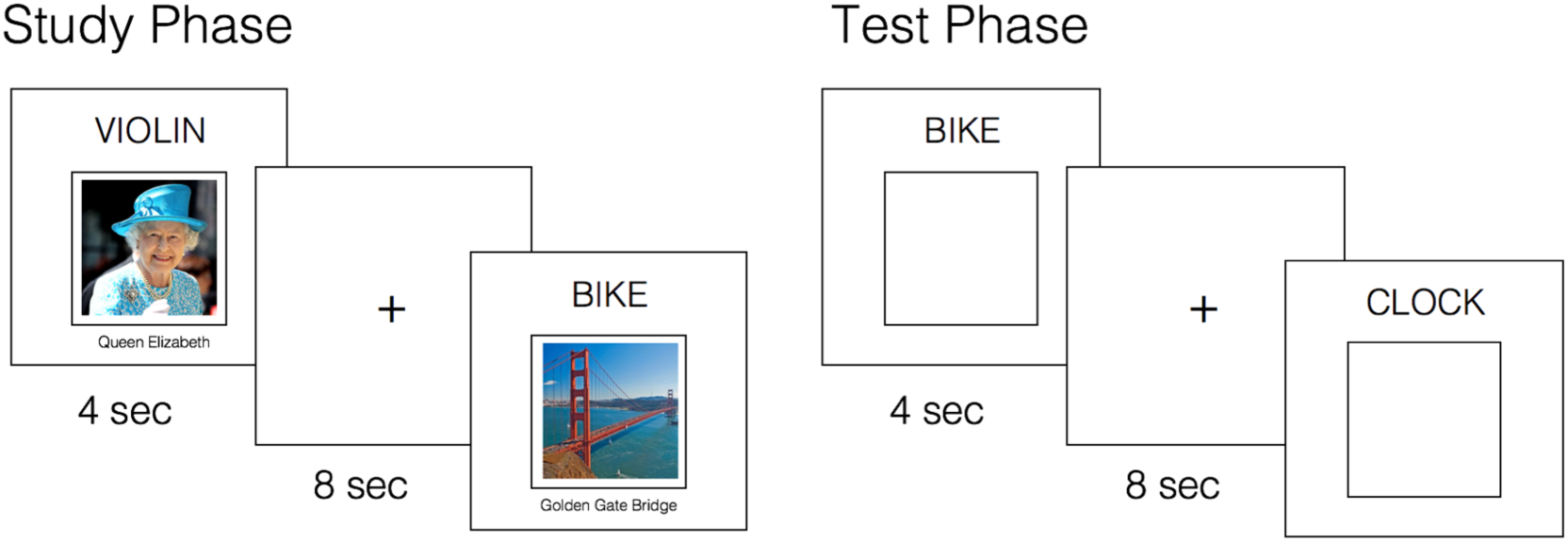
Experimental Paradigm. Concurrent with fMRI, participants intentionally encoded word-picture pairs and completed an associative cued recall test. At test, they were presented with studied words intermixed with novel words, and instructed to recall the associate paired with each word, if old. Participants responded ‘Face’ or ‘Place’ if they could recollect the associated image; ‘Old’ if they recognized the word but could not recollect the associate; ‘New’ if they believed the word was novel. A post-scan cued recall test (not shown, visually identical to the ‘Test Phase’) further probed memory for the specific associate paired with each studied word (see Methods).

**Table 1:** Demographics and Neuropsychological Test Performance

| Measure | Mean (SD) | Range |
| --- | --- | --- |
| Gender | 61 F; 39 M | -- |
| Age (yrs) | 67.96 (5.47) | 60 – 82 |
| Education (yrs) | 16.84 (1.94) | 12 – 20 |
| MMSE | 29.10 (.90) | 26 – 30 |
| CDR | 0 | -- |
| Logical Memory Delayed Recall (/50) | 32.04 (6.16) | 18 – 44 |
| HVLT-R Delayed Recall (/12) | 10.49 (1.68) | 5 – 12 |
| BVMT-R Delayed Recall (/12) | 9.80 (2.16) | 5 – 12 |
| Old/New $d'$ | 2.26 (0.68) | 0.86 – 4.78 |
| Associative $d'$ | 1.64 (0.73) | -0.27 – 3.92 |
| Exemplar-Specific Recall (proportion correct, post-scan) | 0.29 (0.19) | 0.00 – 0.84 |

BVMT-R = Brief Visuospatial Memory Test-Revised; CDR = Clinical Dementia Rating; HVLT-R = Hopkins Verbal Learning Test-Revised; MMSE = Mini Mental State Examination.

To measure pattern completion during retrieval, we used univariate and multivariate analyses focused on a priori regions of interest (ROIs; **Figure 2**). To measure hippocampal function, our primary analyses examined univariate activity in the whole hippocampus bilaterally. In addition, we measured activity in three subfields within the body of the hippocampus –– dentate gyrus/CA3 (DG/CA3), CA1, and subiculum (SUB) –– given prior work suggesting that aging may differentially affect individual hippocampal subfields (39, 41,42) and models predicting differential subfield involvement in pattern completion, including a key role for subfield CA3 (8, 43). To measure cortical reinstatement, we focused on two cortical regions –– ventral temporal cortex (VTC) and angular gyrus (ANG) –– motivated by mounting evidence in healthy younger adults that these two areas support content-rich representations during memory retrieval (10, 25, 44–46), and that their representations may be differentially related to memory-guided behaviour (44–46). Category-level reinstatement (i.e., face/place) was quantified via pattern classification and event-specific reinstatement (e.g., Queen Elizabeth, Golden Gate Bridge) was quantified using encoding-retrieval pattern similarity.

**Figure 2.**
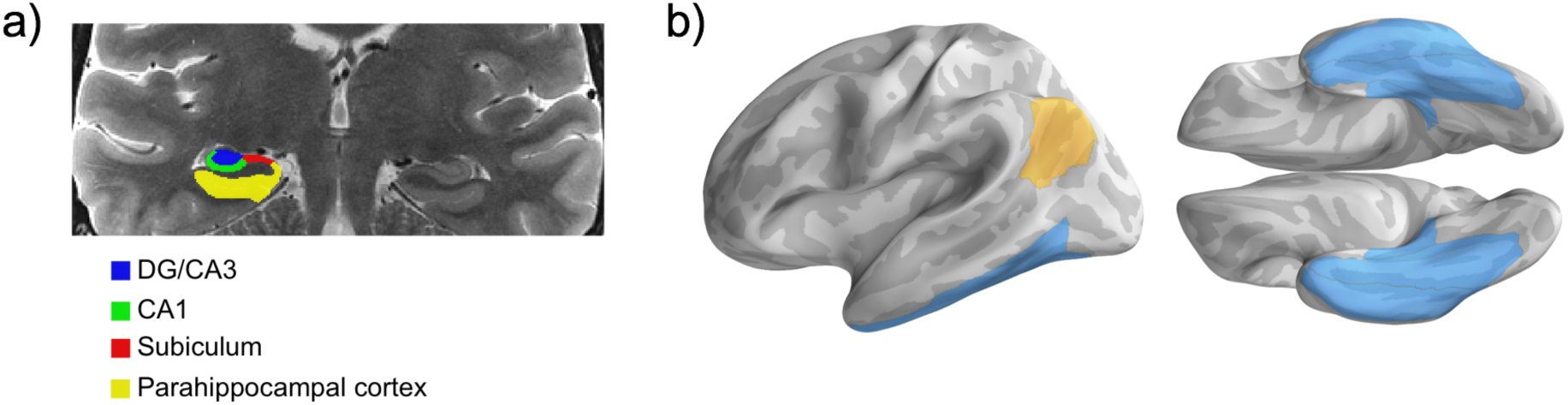
Regions of Interest. (a) Sample MTL subfield demarcations. The whole hippocampus ROI reflects the summation of all subfields (delineated only in the hippocampal body, shown), as well as the hippocampal head and tail (not pictured). (b) Parahippocampal cortex combined with fusiform gyrus and inferior temporal cortex forms the ventral temporal cortex ROI. Ventral temporal cortex (blue) and angular gyrus (gold) masks projected on the fsaverage surface.

## Results

### Behavioural Results

We assessed performance on the associative cued recall task using three measures: 1) old/new *d’* –– discrimination between studied and novel words during the in-scan memory test, irrespective of memory for the associate; 2) associative *d’ ––* correctly remembering the category of associated images encoded with studied words, relative to falsely indicating an associative category to novel words; and 3) post-scan exemplar-specific associative recall – – proportion correct recall of the specific exemplars associated with studied words. Performance on all three measures declined with age (old/new *d*’: *β* = -0.35, *p* < .001; associative *d’*: *β* = -0.30, *p* < .005, **Figure 3a**; post-scan exemplar-specific recall: *β* = -0.34, *p* < .001, **Figure 3b**), but did not vary by sex (*βs* = -0.10, -0.33, -0.23; *p*s *≥* .10) or years of education (*β* = -0.03, -0.02, -0.07; *p*s *>* .47). Critically, despite this decline in performance with age, we also observed considerable variability in performance across individuals in each measure (**Figure 3** and **Table 1**).

**Figure 3.**
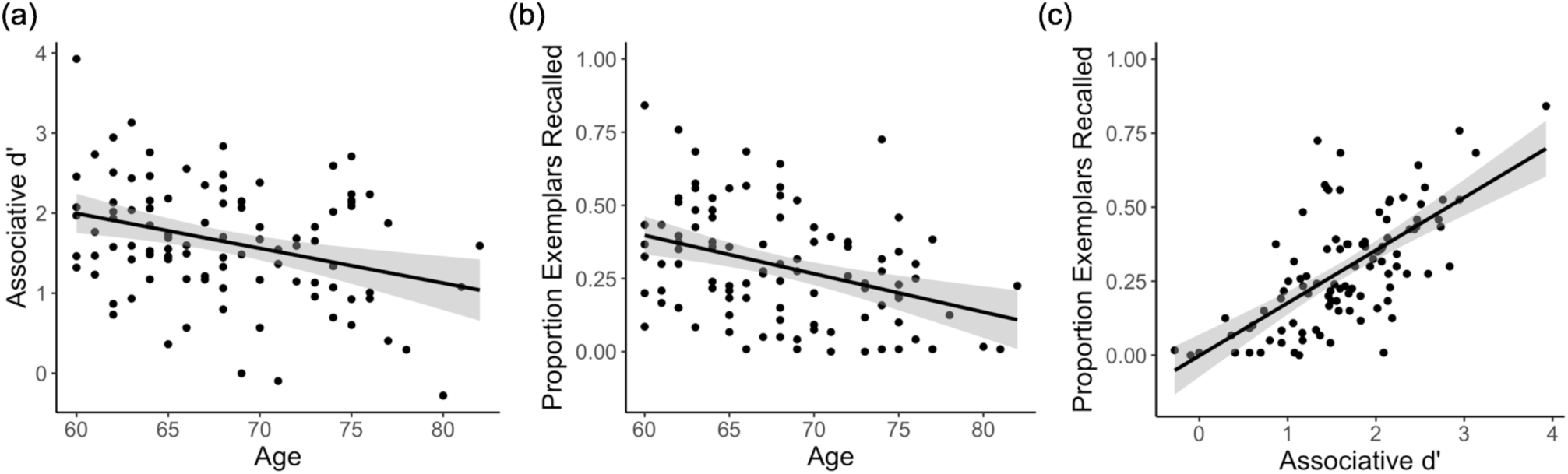
Associative Memory Behavioural Results. (a) In-scanner associative d’ and (b) post-scan exemplar-specific associative recall decline with age. (c) Associative d’ strongly predicts post-scan exemplar-specific associative recall, controlling for the effect of age. Each data point represents a participant; plots show linear model predictions (black line) and 95% confidence intervals (shaded area).

Individual-differences and trial-wise analyses revealed that post-scan associative recall tracked in-scanner associative memory. First, individuals who demonstrated higher associative memory during scanning showed superior recall of the specific exemplars on the post-scan test (controlling for age; *β* = .62, *p* < 10^-12^; **Figure 3c**). Second, trial-wise analysis revealed that making an in-scan associative hit predicted successful post-scan exemplar recall (χ^2^(1) = 159.68, *p* < 10^-36^). These findings suggest that post-scan exemplar-specific retrieval –– while quantitatively lower due to the longer retention interval, change of context, and interference effects –– is a good approximation of recall of the specific exemplar during scanning (relative to simply recalling more general category information).

### fMRI Encoding Classifier Accuracy

Following prior work (e.g., 25, 44-46), cortical reinstatement analyses focused on two a priori ROIs: VTC and ANG. To confirm that activity patterns during word-face and word-place encoding trials were discriminable for each participant in each ROI, we trained and tested a classifier on the encoding data using leave-one-run-out-n-fold cross validation. On average, encoding classifier accuracy was well above chance (50%) using patterns in VTC (M = 98.4%, *p* < .001) and ANG (90.0%, *p* < .001), with classifier accuracy significantly greater in VTC than ANG (*t* (99) = 12.86, *p* < 10^-16^). Classification was above chance in all 100 participants (minimum accuracy of 82.5% (*p* < .001) in VTC and 68.0% (*p* < .005) in ANG). To account for variance in encoding classifier strength (quantified using log odds of the classifier’s probability estimate) on estimates of reinstatement strength during memory retrieval (see **Supplementary Results**, **Figure S1**), we controlled for encoding classifier strength in all subsequent models in which reinstatement strength predicted behavioural variables (memory accuracy, RT), as well as models in which reinstatement strength was the dependent variable (see **Methods** for details).

### Trial-wise Category-level Reinstatement Predicts Memory

We quantified reinstatement of relevant face or scene features (i.e., category-level reinstatement) in VTC and ANG using subject-specific classifiers trained on all encoding phase data for an individual, and tested for cortical reinstatement in the independent retrieval phase data; significance was assessed using permutation testing. Classifier accuracy (**Figure 4a**) was above chance (50%) during associative hits in VTC (M = 68.3%, *p* < .005) and ANG (M = 72.3%, *p* < .001), but did not exceed chance when associative retrieval failed, including on associative miss trials (VTC: 49.8%, *p* = .57; ANG: 50.4%, *p* = .49), item hit (VTC: 53.5%, *p* = .29; ANG: 53.3%, *p* = .31), and item miss trials (VTC: 47.1%, *p* = .68; ANG: 51.6%, *p* = .41; see **Methods** for trial type definitions). Classifier accuracy during associative hits was greater in ANG relative to VTC (*t*(99) = 4.05, *p* < .001). Analyses of the time course of cortical reinstatement during associative hits revealed significant reinstatement effects emerging ∼4-6s post-stimulus onset (**Figure S2**). Analogous category-level reinstatement effects were observed using a pattern similarity approach (i.e., encoding-retrieval similarity (ERS); see **Supplementary Results**).

**Figure 4.**
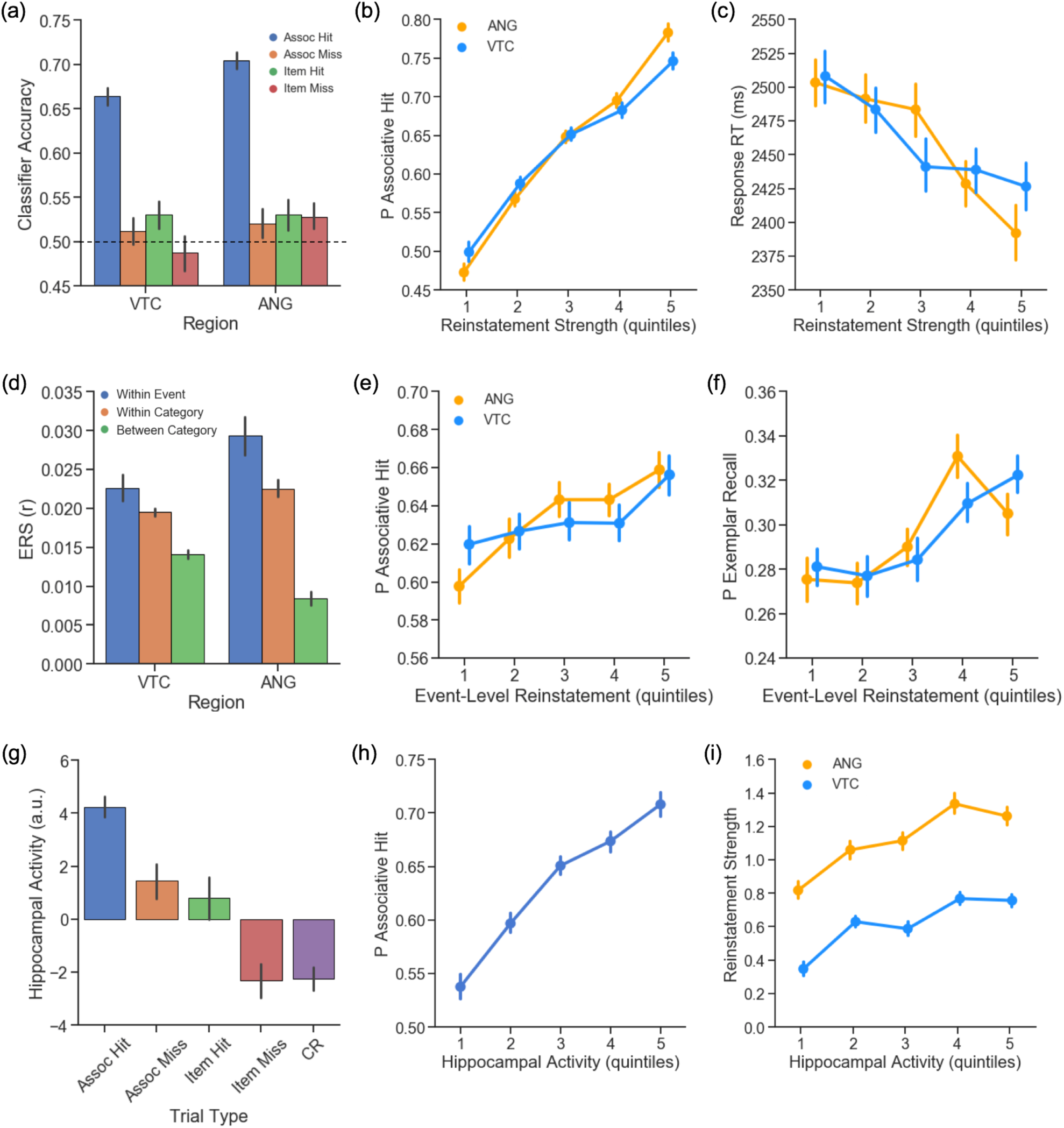
Cortical and Hippocampal Metrics of Pattern Completion during Retrieval. (a) Classifier accuracy is above chance in VTC and ANG during successful, but not unsuccessful, associative retrieval. (b) Trial-wise category reinstatement strength (logits) in VTC and ANG predicts an increased probability of an associative hit and (c) faster decision RT on associative hit trials. (d) Event-level reinstatement (within-event ERS > within-category ERS) is observed during associative hits in VTC and ANG. (e) Trial-wise event-level reinstatement (within-event ERS) predicts the probability of an associative hit and (f) exemplar-specific hit. (g) Hippocampal activity shows a graded response across retrieval conditions. (h) Trial-wise hippocampal activity predicts an increased probability of an associative hit and (i) greater reinstatement strength (logits) in VTC and ANG. For visualization, data for each participant are binned into quintiles based on reinstatement strength (b,c,e,f) and hippocampal activity (h,i). Statistics were conducted on trial-wise data, z-scored within participant. Error bars represent standard error of the mean. VTC = ventral temporal cortex; ANG = angular gyrus. RT = reaction time. ERS = Encoding-Retrieval Similarity.

Evidence for reinstatement during successful, but not unsuccessful, associative retrieval is consistent with theories that posit that reinstatement of event features (here, face or scene features) supports accurate memory-based decisions (here, associate category judgments). More directly supporting this hypothesis, generalized logistic and linear mixed effects models revealed that greater trial-wise cortical reinstatement in VTC and ANG –– quantified using log odds of the classifier’s probability estimate –– predicted (a) an increased probability of an associative hit (VTC: χ^2^(1) = 102.42, *p <* 10^-24^; ANG: χ^2^(1) = 102.42, *p <* 10^-31^; **Figure 4b**), (b) an increased probability of post-scan exemplar-specific recall (VTC: χ^2^(1) = 63.89, *p <* 10^-15^; ANG: χ^2^(1) = 87.44, *p <* 10^-21^; **Figure S3a**), and (c) faster retrieval decision RTs on associative hit trials (VTC: χ^2^(1) = 29.78, *p <* 10^-8^; ANG: χ^2^(1) = 23.39, *p <* 10^-6^; **Figure 4c**). These data provide novel evidence that the strength of category-level reinstatement in VTC and ANG is linked to memory behaviour in cognitively normal older adults (see **Supplementary Results** for analogous ERS findings).

### Trial-wise Event-level Reinstatement Predicts Memory

We next used encoding-retrieval similarity (ERS) to quantify trial-unique, event-specific reinstatement of encoding patterns, comparing the similarity of an event’s encoding and retrieval patterns (within-event ERS) to similarity of encoding patterns from other events from the same category (within-category ERS). Evidence for event-level reinstatement was present in both VTC (*t* (99) = 2.26, *p* < .05) and ANG (*t* (99) = 3.54, *p* < .001) during associative hits (**Figure 4d**). Moreover, the strength of trial-wise event-level reinstatement – – controlling for category-level reinstatement effects (i.e., including within-category ERS as a regressor of noninterest) and univariate activity in each region –– predicted (a) an increased probability of an associative hit (VTC: χ^2^(1) = 1.77, *p =* 0.184; ANG: χ^2^(1) = 7.81, *p <* .005; **Figure 4e**) and (b) an increased probability of post-scan exemplar-specific recall (VTC: χ^2^(1) = 5.33, *p <* .05; ANG: χ^2^(1) = 7.89, *p <* .005; **Figure 4f**), but did not predict decision RT on associative hit trials (VTC: *p* = .837; ANG: *p* = .249). These results demonstrate a relationship between trial-unique, event-specific cortical reinstatement and associative retrieval in older adults.

### Trial-wise Hippocampal Retrieval Activity Predicts Behaviour and Reinstatement

Successful associative retrieval, ostensibly driven by pattern completion, was accompanied by greater hippocampal activity (**Figure 4g**) relative to associative misses (*t*(75) = 4.90, *p* < 10^-6^), item only hits (*t*(59) = 3.87, *p* < .001), item misses (*t*(83) = 8.86, *p* < 10^-13^), and correct rejections (*t*(99) = 11.28, *p* < 10^-16^). Relative to item misses, hippocampal activity was greater during associative misses (*t*(68) = 4.0, *p* < .001) and item only hits (*t*(51) = 5.37, *p* < 10^-6^); activity did not differ between associative misses and item hits (*t* < 1) or between item misses and correct rejections (*t* < 1). Moreover, generalized logistic and linear mixed effects models revealed that greater trial-wise hippocampal activity was linked to (a) an increased probability of an associative hit (χ^2^(1) = 63.23, *p <* 10^-15^; **Figure 4h**), (b) an increased probability of post-scan exemplar-specific recall (χ^2^ (1) = 58.98, *p <* 10^-14^; **Figure S3b**), but (c) not faster associative hit RTs (χ^2^(1) = 2.19, *p =*.139). Thus, the probability of successful pattern-completion-dependent associative retrieval increased with hippocampal activity. This relationship was significant across hippocampal subfields, but greatest in DG/CA3 (see **Supplementary Results** for subfield findings; **Figure S4a**).

Cortical reinstatement is thought to depend on hippocampal pattern completion triggered by retrieval cues (4-7). Consistent with this possibility, the magnitude of trial-wise hippocampal retrieval activity predicted the strength of cortical reinstatement across all retrieval attempts (VTC: χ^2^(1) = 42.38, *p <* 10^-11^; ANG: χ^2^(1) = 34.92, *p <* 10^-19^; **Figure 4i**) and when restricting analyses only to associative hit trials (VTC: χ^2^(1) = 7.01, *p =* .008; ANG: χ^2^(1) = 12.24, *p <* .001). Similarly, hippocampal activity predicted within-event ERS (controlling for within-category ERS) in VTC (all trials: χ^2^(1) = 4.57, *p <* .05; associative hit only: χ^2^(1) =3.87, *p <* .05; see **Figure S5**); this relationship did not reach significance in ANG (all trials: *p =* .388; associative hit only: *p =* .275). Collectively, these results constitute novel evidence for a relationship between trial-wise hippocampal activity and cortical reinstatement in older adults (see **Supplementary Results** for hippocampal subfield findings; **Figure S4b,c**).

### Unique Hippocampal and Cortical Contributions to Associative Retrieval

We next explored whether trial-wise hippocampal activity and trial-wise cortical reinstatement make complementary contributions to associative retrieval success, using nested comparison of logistic mixed effects models. Compared to a model with hippocampal activity, addition of VTC reinstatement strength significantly improved model fit (χ^2^(1) = 103.68, *p* < 10^-24^). Addition of ANG reinstatement to this model further improved model fit (χ^2^(1) = 115.78, *p* < 10^-24^), and all three variables remained significant predictors in the full model (hippocampus: *b =* 0.31, *z =* 8.24, *p <* 10^-16^ ; VTC: *b =* 0.32, *z =* 9.36, *p <* 10^-16^ ; ANG: *b =* 0.52, *z =* 14.42, *p <* 10^-16^). The same approach for exemplar-specific recall similarly revealed that the stepwise addition of reinstatement metrics significantly improved model fit (VTC: χ^2^(1) = 61.17, *p* < 10^-15^; ANG: χ^2^(1) = 65.04, *p* < 10^-16^), with all three variables significant predictors in the full model (hippocampus: *b =* 0.29, *z =* 8.01, *p <* 10^-15^ ; VTC: *b =* 0.21, *z =* 6.44, *p <* 10^-10^ ; ANG: *b =* 0.27, *z =* 9.68, *p <* 10^-16^). Thus, while hippocampal activity predicts cortical reinstatement in VTC and ANG, these three neural responses during retrieval are not redundant predictors of trial-level memory performance. Rather, each makes independent contributions to the probability of a successful associative retrieval decision.

### Effects of Age on Hippocampal and Cortical Indices of Pattern Completion

Our second key aim was to understand how hippocampal pattern completion processes vary across individuals, turning first to the effects of age. To determine whether the trial-wise relationships between our neural metrics and memory behaviour identified in Aim 1 varied as a function of age, we added an interaction term (age*regressor of interest) to each mixed effects model. We observed that age moderated the relationship between reinstatement strength and associative retrieval success in VTC (χ^2^(1) = 6.96, *p <* .01) and marginally in ANG (χ^2^(1) = 3.57, *p* = .059), such that older individuals exhibited a weaker relationship between reinstatement strength and the likelihood of associative retrieval success. In contrast, age did not moderate the relationship between a) hippocampal activity and associative retrieval success (*p* = .643), or b) hippocampal activity and reinstatement strength (VTC: *p* = .777; ANG: *p* = .773). These results suggest that age differentially affects cortical and hippocampal indices of pattern completion, having a particular effect on the translation of cortical evidence to memory behaviour.

To further understand the effects of age, we next asked whether the strength of cortical reinstatement and hippocampal activity during successful associative retrieval (adjusted for relevant nuisance regressors) was reduced with age. Regression analyses revealed that (a) while hippocampal activity during associative hits (associative hit – CR) did not significantly vary with age (*β* = -0.10, *p* = .35; **Figure 5a**), there was (b) an age-related decline in category-level reinstatement strength (i.e., mean logits) during associative hits (VTC: *β* = -0.34, *p* < .0001; ANG: *β* = -0.16, *p* < .05; **Figure 5b-c**), and c) an age-related decline in event-level reinstatement in VTC (*β* = -0.26, *p* < .01; **Figure S6a**), but not ANG (*β* = -0.06, *p* > .55; **Figure S6b**). None of these measures varied with sex or years of education (all *p*s > .24).

**Figure 5.**
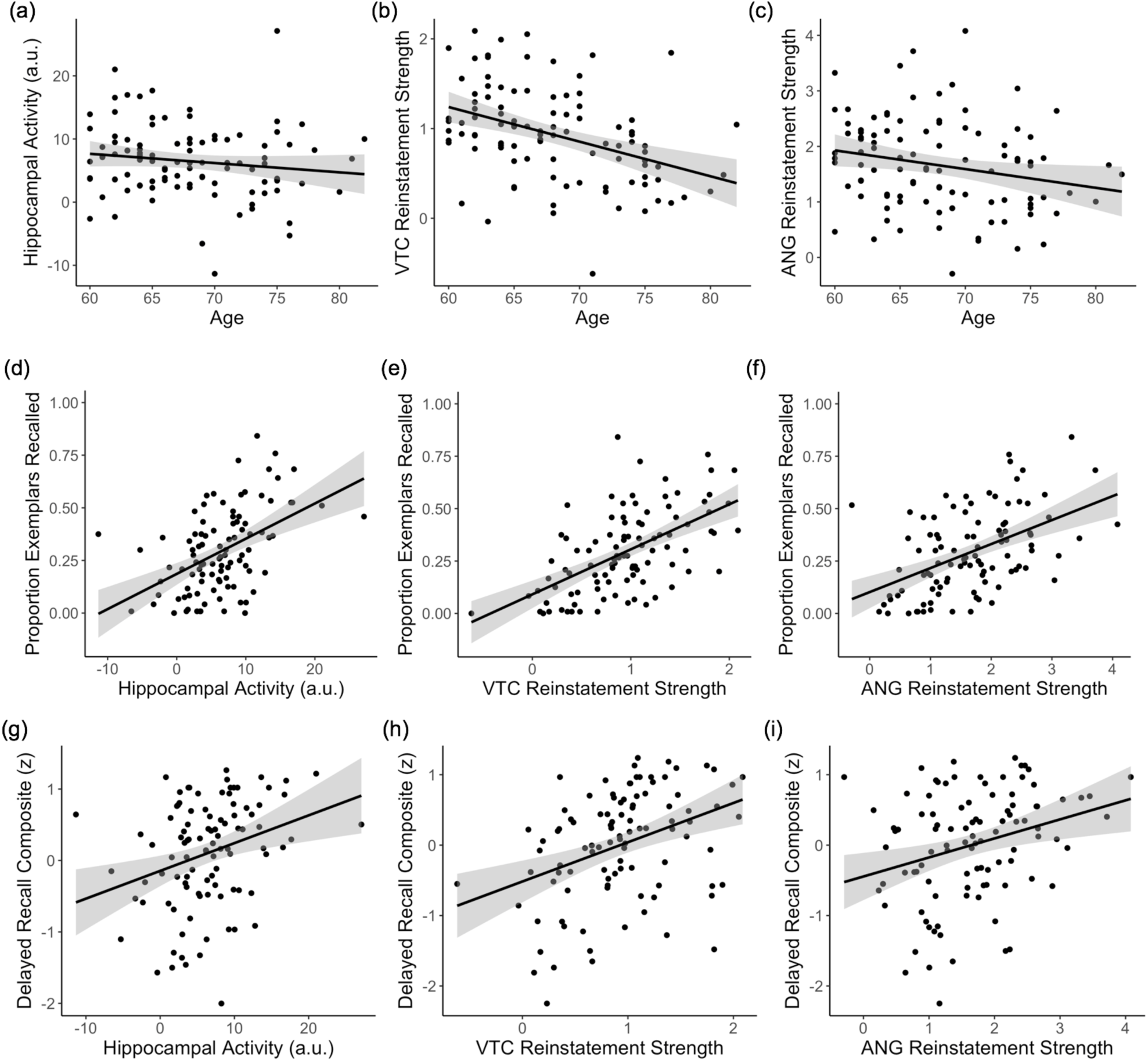
Individual Differences in Pattern Completion Assays. (a-c) Effects of age on hippocampal activity (associative hit – correct rejection) and reinstatement strength (mean logits) in VTC and ANG during associative hits. (d-f) Independent of age, individual differences in hippocampal activity and reinstatement strength in VTC and ANG during associative hits significantly predict exemplar-specific recall. (g-i) Independent of age, individual differences in hippocampal activity and VTC reinstatement strength also explain significant variability in standardized delayed recall performance; the relation with ANG reinstatement did not reach significance. Scatterplots reflect raw values for each measure. See **Supplementary Results (Figure S7)** for partial plots controlling for relevant nuisance variables. Each point represents an individual participant. Plots also show linear model predictions (black line) and 95% confidence intervals (shaded area). VTC = ventral temporal cortex; ANG = angular gyrus.

These cross-sectional age-related declines in category-level and event-level reinstatement during associative hits parallel the age-related decline in associative *d’* and exemplar-specific recall (**Figure 3a-b**). Indeed, age-related change in category-level reinstatement in VTC and (marginally) ANG partially mediated the relationships between age and exemplar-specific recall (VTC: total = -0.37, z = -3.93, *p* < 0.001; direct = -0.19, z = - 2.20, *p* < .05; indirect = -0.17, z = -3.15, *p* < .005, 95% CI = -0.325, -0.093; ANG: total = -0.37, z = -3.93, *p* < 0.001; direct = -0.30, z = -3.40, *p* < .005; indirect = -0.06, z = -1.73, *p* = .08, 95% CI = -0.138, 0.009; see **Supplementary Results** for parallel findings with associative *d’*). Thus, while hippocampal activity during associative hits did not differ by age, cortical reinstatement declined with age and partially mediated the relationship between age and associative memory performance.

### Neural Indices of Pattern Completion Explain Individual Differences in Episodic Memory

We next asked if the strength of neural measures of pattern completion during associative retrieval explain variance in memory performance, independent of age. Separate regression models (adjusted for age and relevant nuisance variables) revealed that individual differences in exemplar-specific recall were predicted by hippocampal activity (*β* = .47, *p* < 10^-7^; **Figure 5d**) and category-level reinstatement strength during associative hits (VTC: *β* = .45, *p* < 10^-6^; ANG: *β* = .52, *p* < 10^-5^, **Figure 5e-f**; see **Figure S7d-f** for partial plots adjusting for nuisance regressors and **Supplementary Results** and **Table S1** for parallel findings with associative *d’*). In contrast, individual differences in event-level reinstatement did not explain significant variance in exemplar-specific recall (all *p*s > .33). Thus, individual differences in the integrity of hippocampal retrieval mechanisms and category-level cortical reinstatement contribute to variability in pattern-completion-dependent (i.e., associative) memory in older adults.

To determine whether these variables explain unique variance in memory performance, we used hierarchical regression (see **Table 2** for model parameters). Compared to a model with age alone (adjusted R^2^ = .126), adding hippocampal activity explained additional variance in performance (model comparison: *F*(1,96) = 29.54, *p* < 10^-7^, adjusted R^2^ = .325). Moreover, adding a single reinstatement metric explained further variance in performance (model comparison: VTC: *F*(1,95) = 22.75, *p* < 10^-6^, adjusted R^2^ = .438; ANG: *F*(1,95) = 8.25, *p* < .01, adjusted R^2^ = .365). However, when VTC and ANG were both included in the same model, reinstatement strength in ANG was no longer a significant predictor (*p* = .412). Thus, in older adults, individual differences in hippocampal activity and cortical reinstatement strength provide complementary information, over and above age, in explaining individual differences in associative memory, whereas indices of reinstatement strength explain shared variance.

**Table 2:** Summary of Regression Analysis Predicting Post-Test Exemplar-Specific Recall

| | Variable | $\beta$ | SE | p | Adjusted $R^2$ |
| --- | --- | --- | --- | --- | --- |
| Step 1 | Age | -0.366 | 0.094 | 0.001*** | 0.126 |
| Step 2 | Age | -0.317 | 0.083 | 0.001*** | 0.325 |
|  | Hippocampal Activity<sub>adj</sub> | 0.472 | 0.087 | 0.001*** |  |
| Step 3a | Age | -0.184 | 0.080 | 0.023* | 0.449 |
|  | Hippocampal Activity<sub>adj</sub> | 0.388 | 0.080 | 0.001*** |  |
|  | VTC Reinstatement<sub>adj</sub> | 0.428 | 0.089 | 0.001*** |  |
| Step 3b | Age | -0.281 | 0.082 | 0.001*** | 0.365 |
|  | Hippocampal Activity<sub>adj</sub> | 0.407 | 0.088 | 0.001**** |  |
|  | ANG Reinstatement<sub>adj</sub> | 0.289 | 0.108 | 0.009** |  |
| Step 4 | Age | -0.184 | 0.080 | 0.023*** | 0.448 |
|  | Hippocampal Activity<sub>adj</sub> | 0.374 | 0.082 | 0.001*** |  |
|  | VTC Reinstatement<sub>adj</sub> | 0.391 | 0.100 | 0.001*** |  |
|  | ANG Reinstatement<sub>adj</sub> | 0.093 | 0.113 | 0.412 |  |
| Step 5 | Age | -0.137 | 0.079 | 0.087~ | 0.485 |
|  | Hippocampal Activity<sub>adj</sub> | 0.335 | 0.080 | 0.001**** |  |
|  | VTC Reinstatement<sub>adj</sub> | 0.377 | 0.089 | 0.001**** |  |
|  | Delayed Recall | 0.299 | 0.110 | 0.008** |  |

*Note.* Adj = Adjusted by relevant nuisance regressors; SE= standard error; VTC = ventral temporal cortex; ANG = angular gyrus; ~ $p < 0.1$ , \* $p < 0.05$ , \*\* $p < .01$ , \*\*\* $p < .001$ \*\*\*\* $p < 10^{-5}$

### Individual Differences in Pattern Completion Predict Independent Measures of Memory

Finally, we examined whether our task-based fMRI measures of pattern completion –– hippocampal activity and cortical reinstatement –– explain individual differences in an independent measure of episodic memory, using a delayed recall composite score collected in a separate neuropsychological testing session (see **Methods**). Controlling for age and sex, hippocampal activity (*β* = 0.19, *p* < .01; **Figure 5g**) and VTC reinstatement strength (*β* = 0.21, *p* < .01; **Figure 5h**) predicted delayed recall score; the relationship with ANG reinstatement strength did not reach significance (*β* = 0.14, *p* = .11; **Figure 5i**; see **Figure S7g-i** for partial plots). Further, as for exemplar-specific recall, we found that hippocampal activity and VTC reinstatement strength explained unique variance in delayed recall performance (hippocampus: *β* = 0.16, *p* < .05; VTC: *β* = 0.20, *p* < .05, adjusted R^2^ = .231).

Given the observed relationships between this standardized neuropsychological measure and the present indices of pattern completion, we asked whether delayed recall score alone could account for the observed relationship between the neural measures and exemplar-specific recall. When delayed recall score was added to the full model (see **Table 2, Step 5**), this measure explained additional variance in exemplar-specific recall (model comparison: *F*(1,94) = 7.45, *p* < .01, adjusted R^2^ = 0.485), but hippocampal activity and VTC reinstatement strength remained significant predictors (hippocampus: *β* = 0.335, *p* < 10^-5^ ; VTC reinstatement: *β* = 0.377, *p* < 10^-5^). Together, these results support the hypothesis that individual differences in the integrity of pattern completion processes, indexed by univariate and pattern-based task-related fMRI metrics, explain variance in memory performance across established hippocampal-dependent assays of episodic memory, and do so in a manner that isn’t captured by simple standardized neuropsychological tests.

## Discussion

Using univariate and multivariate fMRI, the current investigation characterizes the integrity of hippocampal pattern completion during associative retrieval in a large cohort of putatively healthy older adults. We provide novel evidence for unique contributions of hippocampal and cortical indices of pattern completion to a) trial-by-trial differences in episodic remembering in older adults, as well as b) age-related and age-independent individual differences in episodic memory performance. Taken together, these results provide novel insights into the neural mechanisms supporting episodic memory, as well as those driving variability in remembering across older adults.

The present analyses of trial-level brain-behaviour relationships significantly build on work in younger adults (10, 26), demonstrating that trial-wise relationships (a) between hippocampal activity and cortical reinstatement and (b) between each of these neural measures and memory behaviour are present later in the lifespan. While directionality is a difficult to establish with fMRI, these results are consistent with models of episodic retrieval wherein hippocampal pattern completion, triggered by partial cues, drives reinstatement of event representations in the cortex, which supports episodic remembering and memory guided decision making (4–5). Further bolstering this interpretation, the relationship between hippocampal activity and associative retrieval success was qualitatively strongest in DG/CA3 (see **Supplement**), consistent with a key role of CA3 in initiating pattern-completion dependent retrieval (4–8). Moreover, the present results provide novel evidence for stability in the trial-wise relationship between hippocampal activity and (a) cortical reinstatement and (b) associative retrieval success, as neither relationship varied as a function of age.

Consistent with the observed trial-level relationship between hippocampal activity and associative retrieval success, we also demonstrate a positive relationship between the magnitude of hippocampal activity during associative hits and associative memory performance. Our findings complement and build on prior work (36), as we demonstrate that this effect was observed across hippocampal subfields, including DG/CA3, and did not vary significantly as a function of age. These results are compatible with proposals that the relationship between hippocampal ‘recollection success’ effects and memory performance remains stable across the lifespan (36), as well as more broadly, with proposals that preservation of hippocampal function is important for the maintenance of episodic memory in older adults over time (47–48). We note, however, that a negative relationship between hippocampal retrieval activity and memory performance has also been observed in older adults (e.g., 39, 42). Differences across studies may be related to (a) the paradigms and/or contrasts employed (e.g., associative recollection vs. lure discrimination), (b) image resolution (e.g., individual subfields vs. the whole hippocampus), or (c) the make-up of the study population (e.g., cognitively normal or cognitively impaired; 49). Additional well-powered studies of hippocampal retrieval dynamics in older adults are needed to assess the degree to which these variables alter the relationship between hippocampal activity and memory behaviour.

The present results also provide novel insights into the basis of mnemonic decisions in older adults. Specifically, we demonstrate that trial-wise indices of reinstatement strength –– indexed using classifier-derived evidence and encoding-retrieval pattern similarity –– were tightly linked to memory behaviour, including response accuracy and speed. This finding suggests that retrieval was not ‘all or none’, but likely graded (50–52). Indeed, while participants were instructed during scanning to recollect the specific associate, correct category judgments (agnostic to correct exemplar-specific recall) could nonetheless be supported by retrieval of generic category information (i.e., *a place*), prototypical details (e.g., a bridge), specific exemplar details (e.g., the Golden Gate Bridge), or even retrieval of erroneous, but category consistent details (e.g., Niagara Falls). The category-level reinstatement effects observed here likely reflect some combination of these retrieval outcomes, as suggested by the strong correlation between post-scan exemplar-specific recall and within-scan associative d’, along with the observation that the proportion of specific exemplars recalled post-scan was generally lower than correct categorical judgements during scanning (though the former undoubtedly declined due to the longer retention interval and interference effects).

Beyond the strength of reinstatement, the present results cannot adjudicate the nature of the details recalled. For example, both category- and exemplar-specific associative hits could be supported by retrieval of semantic details (e.g., the Golden Gate Bridge), perceptual details (e.g., the bridge was red), or some combination (e.g., vividly recalling the image of the Golden Gate Bridge). One possibility, though speculative, is that VTC and ANG support representations of distinct types of event features (e.g., perceptual features in VTC and semantic features in ANG). This possibility is in line with existing theories (53–54) and also with the present observation that reinstatement strength in VTC and ANG made complementary contributions to retrieval success. Regardless of the precise nature of the details recalled, we demonstrate that, as in younger adults (10, 44, 51), recovery of stronger mnemonic evidence was associated with greater accuracy and faster responses, and this was true for representations supported by VTC and ANG alike. This relationship may reflect reduced demands on post-retrieval monitoring and selection processes and/or greater confidence in the face of stronger mnemonic evidence. Interestingly, the strength of the trial-level relationship between reinstatement strength and behaviour weakened with increased age. This could be related to age-related changes in decision criteria, retrieval monitoring ability, response strategies, or some combination of these factors. Future work is needed to explore the specific neurocognitive basis of this intriguing effect, which likely involves interactions between the medial temporal lobe and frontoparietal regions (26, 55).

Although we observed robust group-level cortical reinstatement effects during associative hits, reinstatement strength declined with age, and partially mediated the relationship between age and episodic memory. These data provide neuroimaging evidence in support of proposals that age-related episodic memory decline is driven, in part, by a loss of specificity or precision in mnemonic representations, a possibility that has been well-supported by behavioural evidence (56–58). Importantly, the effect of age on reinstatement strength, and the relationship between reinstatement strength and memory performance, was observed after accounting for variance in encoding classifier performance, a putative assay of cortical differentiation (i.e., the ability to establish distinct neural patterns associated with different visual stimulus categories) during memory encoding. Thus, although we found that encoding classifier strength was a strong predictor of reinstatement strength, consistent with prior work (35) and existing proposals regarding dedifferentiation of cortical representations in older adults (59–61), the present results suggest that the observed variance in reinstatement strength does not simply reflect downstream effects of cortical differentiation. Instead, variance in reinstatement strength likely also provides information about the precision with which event representations are retrieved in older adults.

Interestingly, while cortical reinstatement is a putative read-out of pattern completion, a possibility further supported by the present data, the hippocampal and cortical metrics defined here explained unique variance in memory performance, both at the trial level and across individuals. Indeed, these measures together explained nearly three times as much variance in exemplar-specific associative recall as age alone (**Table 2**). One possibility is that hippocampal activity and cortical reinstatement strength index distinct aspects of recollection: retrieval success vs. retrieval precision, respectively (e.g., 52, 62). That is, whereas increases in hippocampal activity may signal recollection of some event details, this signal alone may not indicate the fidelity or precision with which the event is recollected. Conversely, reinstatement strength likely provides more information about the contents of recollection, including the specificity or precision of mnemonic representations (e.g., recall of generic as opposed to exemplar-specific details), and perhaps even the nature of the details recollected (i.e., perceptual vs semantic). Alternatively, representations reinstated in cortex may be differentially affected by top-down goal representations and decision processes (44–45, 55), which contribute unique variance in memory performance beyond that explained by hippocampal-cortical event replay. Future work is needed to examine whether the unique variance explained by cortical reinstatement relates to frontoparietal control and decision processes in older adults.

Indeed, it is important to note that variability in episodic remembering, and indeed variability in the strength of the present pattern completion metrics, is likely influenced by a number of variables, only some of which are measured here. For example, aging may affect other processes at retrieval, including elaboration of retrieval cues (63) and post-retrieval monitoring and selection (61, 64), as well as factors at encoding, including the differentiation of stimulus representations (59–61), goal-directed or sustained attention (65–66), and elaborative or ‘strategic’ encoding processes (67–68). These variables could vary both within individuals (i.e., across trials), as well as between individuals (e.g., trait level differences). The manner in which these variables impact pattern completion processes at retrieval, or make independent contributions to episodic remembering in older adults, is an important direction for future work. Nevertheless, the present results provide compelling initial evidence that (a) hippocampal and cortical indices of pattern completion play a central role in determining whether individual events will be remembered or forgotten, (b) that predicted relationships between hippocampal activity, reinstatement strength, and associative memory retrieval can be observed even late in the lifespan, and (c) and that these neural metrics explain unique variance in memory performance across individuals.

Hippocampal and cortical indices of pattern completion not only explained variance in our primary associative memory measures, but also in delayed recall performance on standardized neuropsychological tests –– among the most widely used assays of episodic memory in the study of aging and disease. The relationship between these measures, collected during separate testing sessions, suggests that the neural indices derived from task-based fMRI are tapping into stable individual differences, and may represent a sensitive biomarker of hippocampal and cortical function. Critically, we also demonstrate that these neural and neuropsychological test measures explained unique variance in associative memory, together accounting for 50% of the variance in exemplar-specific recall across individuals. This not only indicates that the present neural indices provide information that cannot be garnered from paper and pencil tests alone, but also suggests that we can combine these neural metrics with existing measurement tools to build more accurate models to explain individual differences in memory performance in older adults. An important direction for future work is to assess whether combining task-related neural measures, such as those identified here, with other known biomarkers of brain health and disease risk (e.g., in vivo measures of amyloid and tau accumulation, hippocampal volume, white matter integrity; 69-70) can further increase sensitivity for explaining individual differences in memory performance, as well as predicting future disease risk and memory decline prior to the emergence of clinical impairment.

Taken together, the present results significantly advance our understanding of fundamental retrieval processes supporting episodic memory in cognitively normal older adults. By exploring how neural indices of pattern completion vary –– both across trials and across individuals –– these findings demonstrate that hippocampal activity and cortical reinstatement during memory retrieval provide a partial account for why and when older adults remember, and they predict which older adults will perform better than others across multiple widely adopted assays of episodic memory. Our findings also underscore the striking heterogeneity in brain and behaviour among cognitively normal older adults, and lend support to the hypothesis that this high within-group variance likely contributes to the wealth of mixed findings in the literature, particularly for traditional group-level comparisons in the context of small-to-moderate sample sizes. Collectively, our findings illustrate how an individual differences approach can advance understanding of the neurocognitive mechanisms underlying when and which older adults are more likely to remember.

## Methods

### Participants

One hundred and five cognitively healthy older adults (aged 60-82 yrs; 65 female) participated as part of the Stanford Aging and Memory Study. Eligibility included: normal or corrected-to-normal vision and hearing; right-handed; native English speaking; no history of neurological or psychiatric disease; a Clinical Dementia Rating score of zero (CDR; 71); and performance within the normal range on a standardized neuropsychological assessment (see Neuropsychological Testing). Data collection spanned multiple visits: Neuropsychological assessment was completed on the first visit and the fMRI session occurred on the second visit, with the exception of nine participants who completed the fMRI session on the same day as the neuropsychological testing session. Visits took place ∼6.18 weeks apart on average (range = 1–96 days). Participants were compensated $50 for the clinical assessment and $80 for the fMRI session. All participants provided informed consent in accordance with a protocol approved by the Stanford Institutional Review Board. Data from five participants were excluded from all analyses due to excess head motion during scanning (see fMRI Pre-processing), yielding a final sample of 100 older adults (60-82 yrs; 61 female; see Table 1 for demographics).

### Neuropsychological Testing

Participants completed a neuropsychological test battery consisting of standardized tests assessing a range of cognitive functions, including episodic memory, executive function, visuospatial processing, language, and attention. Scores were first reviewed by a team of neurologists and neuropsychologists to evaluate cognition and reach a consensus assessment that each participant was cognitively healthy, defined as performance on each task within 1.5 standard deviations of demographically adjusted means. Subsequently, a composite delayed recall score was computed for each participant by (a) z-scoring the delayed recall subtest scores from the Logical Memory (LM) subtest of the Wechsler Memory Scale, 3rd edition (WMS-III; 72), Hopkins Verbal Learning Test-Revised (HVLT-R; 73), and the Brief Visuospatial Memory Test-Revised (BVMT-R; 74), and (b) then averaging. This composite score declined with age (β = -0.21, p < .005), was lower in males than females (β = -0.35, p < .05), but did not vary with years of education (β = 0.07, p > .31).

### Materials

Stimuli comprised words paired with colour photos of faces and scenes obtained from online sources. For each participant, 120 words (out of 150 words total) were randomly selected and paired with the pictures (60 word-face; 60 word-place) during a study phase, and these 120 words plus the remaining 30 words (foils) appeared as cues during the retrieval phase. Words were concrete nouns (e.g., “banana”, “violin”) between 4 and 8 letters in length. Faces corresponded to famous people (e.g., “Meryl Streep”, “Ronald Reagan”) and included male and female actors, musicians, politicians, and scientists. Places corresponded to well-known locations (e.g., “Golden Gate Bridge”, “Niagara Falls”) and included manmade structures and natural landscapes from a combination of domestic and international locations.

### Behavioural Procedure

Prior to scanning, participants completed a practice session that comprised an abbreviated version of the task (12 word-picture pairs not included in the scan session). This ensured that participants understood the task instructions and were comfortable with the button responses. Participants had the option to repeat the practice round multiple times if needed to grasp the instructions.

Next, concurrent with fMRI, participants performed an associative memory task consisting of five rounds of alternating encoding and retrieval blocks (Figure 1). In each encoding block, participants viewed 24 word-picture pairs (12 word-face and 12 word-place) and were asked to intentionally form an association between each word and picture pair. To ensure attention to the pairs, participants were instructed to indicate via button press whether they were able to successfully form an association between items in the pair. Following each encoding block, participants performed a retrieval task that probed item recognition and associative recollection. In each block, 24 target words were interspersed with 6 novel (foil) words; participants made a 4-way memory decision for each word. Specifically, if they recognized the word and recollected the associated image, they responded either ‘Face’ or ‘Place’ to indicate the category of the remembered image; if they recognized the word but could not recollect sufficient details to categorize the associated image, they responded ‘Old’; if they did not recognize the word as studied, they responded ‘New’. Responses were made via right-handed button presses, with four different finger assignments to the response options counterbalanced across participants. Using MATLAB Psychophysics Toolbox (75), visual stimuli were projected onto a screen and viewed through a mirror; responses were collected through a magnet-compatible button box.

During both encoding and retrieval blocks, stimuli were presented for 4s, followed by an 8-s inter-trial fixation. During retrieval blocks, the probe word changed from black to green text when there was 1s remaining, indicating that the end of the trial was approaching and signaling participants to respond (if they had not done so already). After the MR scan session, a final overt cued-recall test was conducted outside the scanner to evaluate the degree to which participants were able to recollect the specific face or place associated with each target word. On this post-test, participants were presented with studied words, in random order, and asked to provide the name of the associate or, if not possible, a description of the associate in as much detail as they could remember. The post-test was self-paced, with responses typed out on a keyboard; participants were instructed to provide no response if no details of the associate could be remembered.

### Memory Response Classification

The fMRI retrieval trials were classified into six conditions: associative hits (AH; studied words for which the participant indicated the correct associate category), associative misses (AM; studied words for which the participant indicated the incorrect associate category), item hits (IH; studied words correctly identified as ‘old’), item misses (IM; studied words incorrectly identified as ‘new’), item false alarms (FAI; foils incorrectly called ‘old’), associative false alarms (FAA; foils incorrectly indicated as associated with a ‘face’ or a ‘place’), and correct rejections (CR; foils correctly identified as ‘new’). Because the number of false alarms was low (M = 5.1, SD = 4.7), these trials were not submitted to fMRI analysis.

In-scanner associative memory performance was estimated using a discrimination index, associative d’. Hit rate was defined as the rate of correct category responses to studied words (AH) and the false alarm rate was defined as the rate of incorrect associative responses to novel words (FAA). Thus, associative d’ = Z(‘AH’ | OLD / All OLD) – Z(‘FAA’ | NEW / All NEW). We additionally calculated an old/new discrimination index to assess basic understanding of and ability to perform the task. Here, hit rate was defined as the rate of correct old responses to studied words, irrespective of associative memory (AH, AM, IH), and the false alarm rate was defined as the rate of incorrect old responses to novel words (FAA, FAI). Thus, old/new d’ = Z(‘AH’ + ‘AM’ + ‘IH’ | OLD / All OLD) – Z(‘FAA’ + ‘FAI’ | NEW / All NEW).

The post-test data were analysed using a semi-automated method. Participants’ typed responses were first processed with in house R code to identify exact matches to the name of the studied image. Responses that did not include exact matches were flagged, and subsequently assessed by a human rater, who determined the correspondence between the description provided by the participant and the correct associate. We computed the proportion of studied words for which the associate was correctly recalled (Exemplar Correct/All Old). One participant did not complete the post-test, leaving 99 participants in all analyses of the post-test data.

### MRI Data Acquisition

Data were acquired on a 3T GE Discovery MR750 MRI scanner (GE Healthcare) using a 32-channel radiofrequency receive-only head coil (Nova Medical). Functional data were acquired using a multiband EPI sequence (acceleration factor = 3) consisting of 63 oblique axial slices parallel to the long axis of the hippocampus (TR = 2 s, TE = 30 ms, FoV = 215 mm x 215 mm, flip angle = 74, voxel size = 1.8 × 1.8 × 2 mm). To correct for B0 field distortions, we collected two B0 field maps before every functional run, one in each phase encoding direction. Two structural scans were acquired: a whole-brain high-resolution T1-weighted anatomical volume (TR = 7.26 ms, FoV = 230 mm × 230 mm, voxel size = 0.9 x 0.9 x 0.9 mm, slices = 186), and a T2-weighted high-resolution anatomical volume perpendicular to the long axis of the hippocampus (TR = 4.2 s, TE = 65 ms, FOV = 220 mm, voxel size = 0.43 × 0.43 × 2 mm; slices = 29). The latter was used for manual segmentation of hippocampal subfields and surrounding cortical regions (76).

### fMRI Pre-processing

Data were processed using a workflow of FSL (77) and Freesurfer (78) tools implemented in Nipype (79). Each timeseries was first realigned to its middle volume using normalized correlation optimization and cubic spline interpolation. To correct for differences in slice acquisition times, data were temporally resampled to the TR midpoint using sinc interpolation. Finally, the timeseries data were high-pass filtered with a Gaussian running-line filter using a cutoff of 128 s. The hemodynamic response for each trial was estimated by first removing the effects of motion, trial artefacts (see **Supplementary Methods**), and session from the timeseries using a general linear model. The residualized timeseries was then reduced to a single volume for each trial by averaging across TRs 3-5 (representing 4-10s post-stimulus onset), corresponding to the peak of the hemodynamic response function. To preserve the high resolution of the acquired data, the data were left unsmoothed.

Images with motion or intensity artifacts were automatically identified as those TRs in which total displacement relative to the previous frame exceeded 0.5mm or in which the average intensity across the whole brain deviated from the run mean by greater than five standard deviations. Runs in which the number of artifacts identified exceeded 25% of timepoints, as well as runs in which framewise displacement exceeded 2mm, were excluded. These criteria led to exclusion of data from five participants who exhibited excess head motion across runs, as well as exclusion of one study and test run from an additional participant. Across all included runs from 100 participants, an average of 2.4 (SD = 3.7) encoding phase volumes (1.7% of volumes) and 2.6 (SD = 4.2) retrieval phase volumes (1.5% of volumes) were identified as containing an artifact. Trials containing fMRI artifacts were excluded from all analyses. To control for potential residual effects of head motion on our primary variables of interest, we adjusted each variable of interest by mean framewise displacement using linear regression (see **Supplementary Results**).

Using Freesurfer, we segmented the T1-weighted anatomical volume at the gray-white matter boundary and constructed tessellated meshes representing the cortical surface (78). Functional data from each run were registered to the anatomical volume with a six degrees-of-freedom rigid alignment optimizing a boundary-based cost function (80). Finally, runs 2–4 were resampled into the space of run 1 using cubic spline interpolation to bring the data into a common alignment. All analyses were thus performed in participant native space, avoiding normalization to a group template.

### Regions of Interest

Our analyses focus specifically on hippocampal pattern completion processes –– via hippocampal univariate activity and multivariate cortical reinstatement metrics –– in the aging brain. Thus, analyses were conducted in three a priori regions of interest (ROI), selected based on existing theoretical and empirical work to optimize the measurement of this process. Analyses of task-evoked univariate activity were focused on the hippocampus, whereas multivoxel pattern analyses were conducted in ventral temporal cortex (VTC) and angular gyrus (ANG), two cortical areas that have been reliably linked to cortical reinstatement in healthy younger adults (10, 25, 44–46). All ROIs were bilateral and defined in participants’ native space (**Figure 2**).

The hippocampal mask was defined manually using each participant’s high-resolution T2-weighted structural image using established procedures (76), and comprised the whole hippocampus (see **Supplementary Results** for analysis of hippocampal subfields). The VTC mask was composed of three anatomical regions: parahippocampal cortex, fusiform gyrus, and inferior temporal cortex. The fusiform gyrus and inferior temporal cortex masks were generated from each participant’s Freesurfer autosegmentation volume using bilateral inferior temporal cortex and fusiform gyrus labels. These were combined with a manually defined bilateral parahippocampal cortex ROI, defined using established procedures (76), to form the VTC mask. The ANG ROI was defined by the intersection of the Freesurfer inferior parietal lobe label and the Default Network of the Yeo 7 network atlas (81), defined on the Freesurfer average (fsaverage) cortical surface mesh. This intersection was used to confine the ROI to the inferior parietal nodes of the Default Mode Network, which predominantly encompasses ANG (45). To generate ROIs in participants’ native space from the fsaverage space label, we used the approach detailed in Waskom and colleagues (55), which uses the spherical registration parameters to reverse-normalize the labels, and then converts the vertex coordinates of labels on the native surface into the space of each participant’s first run using the inverse of the functional to anatomical registration. Participant-specific ROIs were then defined as all voxels intersecting the midpoint between the gray-white and gray-pial boundaries.

### Multivoxel Pattern Classification

Our primary measure of cortical reinstatement during memory retrieval was derived from multivoxel classification analysis. Classification was implemented using Scikit-learn (82), nilearn (83), nibabel (84), and in house Python scripts, and performed using L2-penalized logistic regression models as instantiated in the LIBLINEAR classification library (regularization parameter C =1). These models were fit to preprocessed BOLD data from VTC and ANG that were reduced to a single volume for each trial by averaging across TRs 3-5. Prior to classification, the sample by voxel matrices for each region were scaled across samples within each run, such that each voxel had zero mean and unit variance. A feature selection step was also conducted, in which a subject-specific univariate contrast was used to identify the top 250 voxels that were most sensitive to each category (face, place) during encoding, yielding a set of 500 voxels over which classification analyses were performed. Prior to each of 10 iterations of classifier training, the data were subsampled to ensure an equal number of face and scene trials following exclusion of trials with artefacts.

To first validate that classification of stimulus category (face/place) during encoding was above chance for each ROI, we used a leave-one-run-out n-fold cross-validation procedure on the encoding data. This yielded a value of probabilistic classifier output for each trial, representing the degree to which the encoding pattern for a trial resembled the pattern associated with a face or place trial. This output was converted to binary classification accuracy indicating whether or not a given test trial was correctly classified according to the category of the studied picture. Here we report the average classifier accuracy across folds for each participant in each ROI.

To measure cortical reinstatement during memory retrieval, we trained a new classifier on all encoding phase data, and then tested on all retrieval phase data. For each retrieval trial, the value of probabilistic classifier output represented a continuous measure of the probability (range 0-1) that the classifier assigned to the relevant category for each trial (0 = certain place classification, 1 = certain face classification). For assessment of classifier performance across conditions (associative hits, associative misses, item only hits, and item misses) and ROI (VTC, ANG), we converted this continuous measure of classifier evidence to binary classification accuracy, indicating whether or not a given retrieval trial was correctly classified according to the category of the studied picture.

The significance of classifier performance for each condition and ROI was assessed using permutation testing. We generated a null distribution for each participant by shuffling the trial labels over 1000 iterations for each of the 10 subsampling iterations, calculating mean classifier accuracy for each iteration. We then calculated the mean number of times the permuted classifier accuracy met or exceeded observed classifier accuracy to derive a p value indicating the probability that the observed classifier accuracy could arise by chance.

For trial-wise analyses relating cortical reinstatement strength to memory behaviour (e.g., associative retrieval accuracy and reaction time) and other neural variables (e.g., hippocampal BOLD), a continuous measure of reinstatement strength was derived by calculating the logits (log odds) of the probabilistic classifier output on each trial. Reinstatement strength was signed in the direction of the correct associate for a given trial, such that, regardless of whether the trial was a face or place trial, the evidence was positive when the classifier guessed correctly, and negative when the classifier guessed incorrectly. The magnitude of reinstatement strength was thus neutral with respect to which associate category (face or place) was retrieved.

### Pattern Similarity Analysis

To complement the classification analyses, we used pattern similarity analyses to measure cortical reinstatement. This approach involved computing the similarity (Pearson correlation) between trial-wise activity patterns extracted from ROIs during encoding and retrieval (i.e., encoding-retrieval similarity; ERS). This analysis approach affords the opportunity to not only examine reinstatement at the categorical level (i.e., within-category ERS – between-category ERS) but also at the trial-unique item level (i.e., within-event ERS – within-category ERS). For this analysis, we again used the voxelwise activity patterns for each ROI, computing the correlation between encoding and retrieval patterns separately for successful (i.e., associative hits) and unsuccessful (i.e., associative misses, item only hits, item misses) retrieval trials, such that the events being compared (within-event, within-category, between-category) were matched on associative retrieval success. All correlations were Fisher transformed before computing the mean correlation between different events of interest.

### Statistical Analysis

All statistical analyses were implemented in the R environment (version 3.4.4). Trial-wise analyses were conducted using mixed effects models (linear and logistic) using the lmer4 statistical package (85). Each model contained fixed effects of interest, a random intercept modeling the mean subject-specific outcome value, and a random slope term modeling the subject-specific effect of the independent variable of interest (e.g., hippocampal activity, reinstatement strength). Models also contained relevant nuisance regressors, including stimulus category, ROI encoding classifier strength (when reinstatement strength (logits) was the independent or dependent variable), ROI univariate activity in category-selective voxels (when reinstatement strength (logits) was the independent variable or dependent variable); the significance of these variables was explored in separate models (see **Supplementary Results**). Random slopes were uncorrelated from random intercepts to facilitate model convergence. The significance of effects within mixed-model regressions was obtained using log-likelihood ratio tests, resulting in χ^2^ values and corresponding p-values. A Wald z-statistic was additionally computed for model parameters to determine simultaneous significance of coefficients within a given model. All continuous variables were z-scored within participant across all trials prior to analysis.

Individual differences analyses were conducted using multiple linear regression. In all regression models, each neural variable was adjusted by the relevant nuisance regressors, namely head motion (mean framewise displacement) and, where relevant, ROI encoding classifier strength (mean logits). Age-independent models adjusted memory scores by age. Main text figures depict raw values for interpretability (see **Supplementary Figures** for partial plots). Hierarchical Regression was used to assess the relative contributions of each independent variable to memory performance. F ratio statistics were used to determine change in explained variance (R^2^) at each step compared to the previous step. The explanatory power of each regression model was evaluated descriptively using the explained variance (adjusted R^2^). All continuous variables were z-scored across participants prior to analysis, producing standardized coefficients. All analyses used a two-tailed level of 0.05 for defining statistical significance.

## Supporting information

Supplementary Results

## Acknowledgements

The Stanford Aging and Memory Study (SAMS) is supported by the National Institute on Aging (R01AG048076, R21AG058111, and R21AG058859), Stanford’s Center for Precision Health and Integrated Diagnostics (PHIND), Stanford’s Wu Tsai Neurosciences Institute. We are grateful to colleagues Adam Kerr, Hua Wu, Michael Perry, and Laima Baltusis at the Stanford’s Center for Cognitive and Neurobiological Imaging (CNI) for their assistance in fMRI data acquisition, Clementine Chou, Madison Kist, and Austin Salcedo for their assistance with data collection, and the SAMS volunteers for their participation in the study. The current institutions for the following authors are as follows: AN – University of Texas at Austin; CAF – Yale University; CPL – Johns Hopkins University; JDB – UC San Diego Medical School; MJ – Columbia University; MKT – Columbia University; VAC – San Jose State University; WG – University of Oregon.

