## Supplementary Results for "Hippocampal and cortical mechanisms at retrieval explain variability in episodic remembering in older adults"

**Supplementary Materials**

### Main Text Supplementary Figures

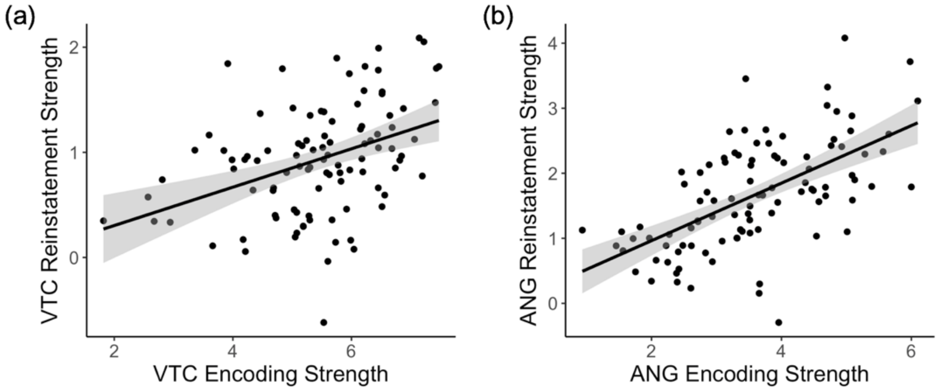


*Figure S1. Relationship between Encoding Strength (logits) and Reinstatement Strength (logits) in (a) VTC and (b) ANG. In both cases, encoding strength was a significant predictor of reinstatement strength. Each point on the scatterplot represents an individual subject. Plots also show the linear model predictions (black line) and 95% confidence interval (shaded area). VTC = ventral temporal cortex; ANG = angular gyrus.*

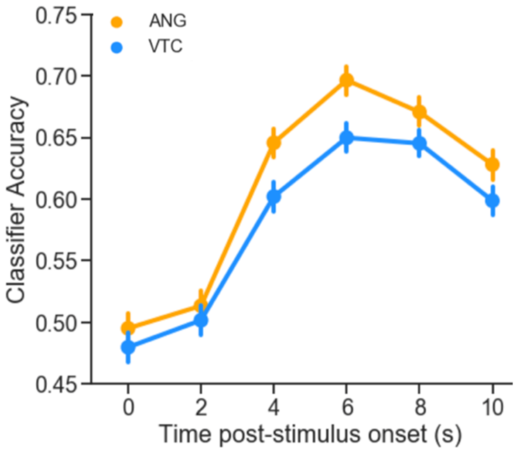


*Figure S2. Time Course of Cortical Reinstatement during Associative Hits. Reinstatement effects emerged at approximately 4-6s post-stimulus onset in both VTC and ANG. Error bars represent standard error of the mean. VTC = ventral temporal cortex; ANG = angular gyrus.*

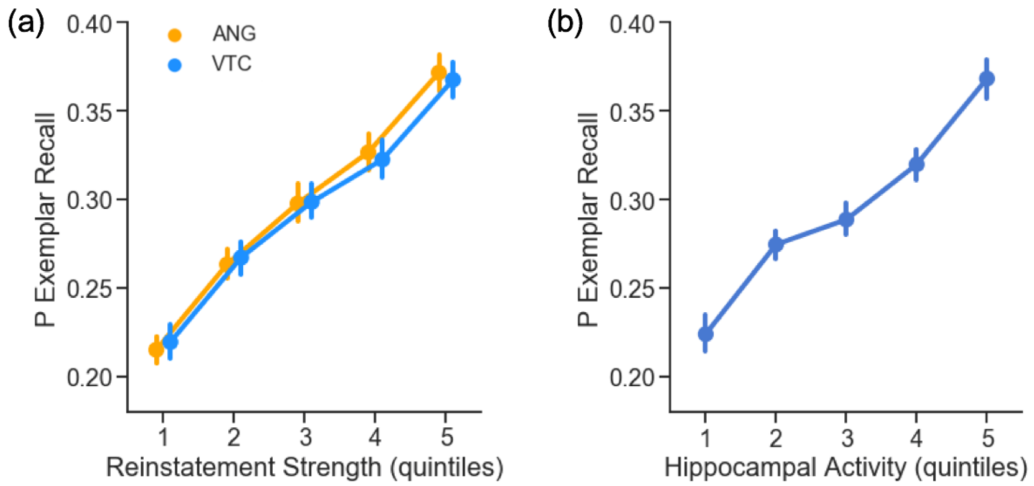


*Figure S3. In-scanner Pattern Completion Metrics Predict Post-Scan Exemplar-Specific Recall. Trial-wise estimates of (a) category reinstatement (logits) and (b) hippocampal activity predict an increased probability of exemplar-specific recall in the post-scan memory test. For visualization, data for each participant were binned into quintiles based on reinstatement or hippocampal activity. Statistics were conducted on trial-wise data, z-scored within participant. Error bars represent standard error of the mean. VTC = ventral temporal cortex; ANG = angular gyrus.*

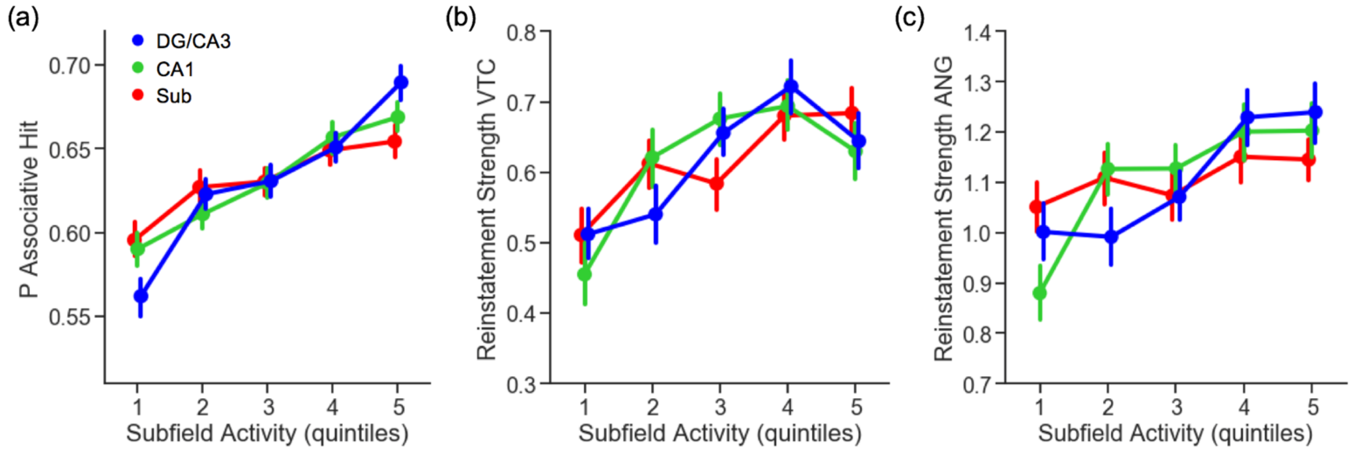


*Figure S4: Hippocampal Subfield Activity during Associative Retrieval. (a) Activity in DG/CA3, CA1, and Sub predicts associative retrieval success, (b) category reinstatement strength in VTC, and (c) category reinstatement strength in ANG. For visualization, data for each participant were binned into quintiles based on hippocampal subfield activity. Statistics were conducted on trial-wise data, z-scored within participant. Error bars represent standard error of the mean. VTC = ventral temporal cortex; ANG = angular gyrus; DG = Dentate Gyrus; Sub = Subiculum.*

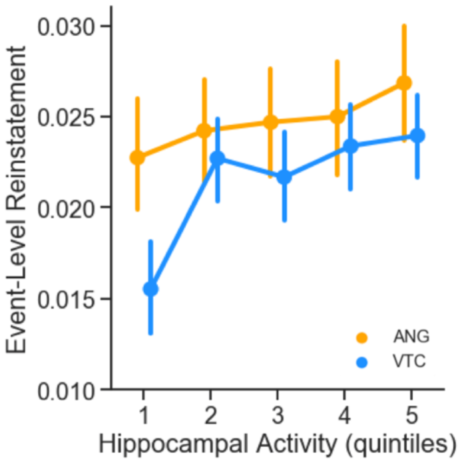


*Figure S5. Trial-wise Hippocampal Activity Predicts Within-Event ERS in VTC. For visualization, data for each participant were binned into quintiles based on hippocampal activity. Statistics were conducted on trial-wise data, z-scored within participant. Error bars represent standard error of the mean. VTC = ventral temporal cortex; ANG = angular gyrus. ERS = Encoding Retrieval Similarity.*

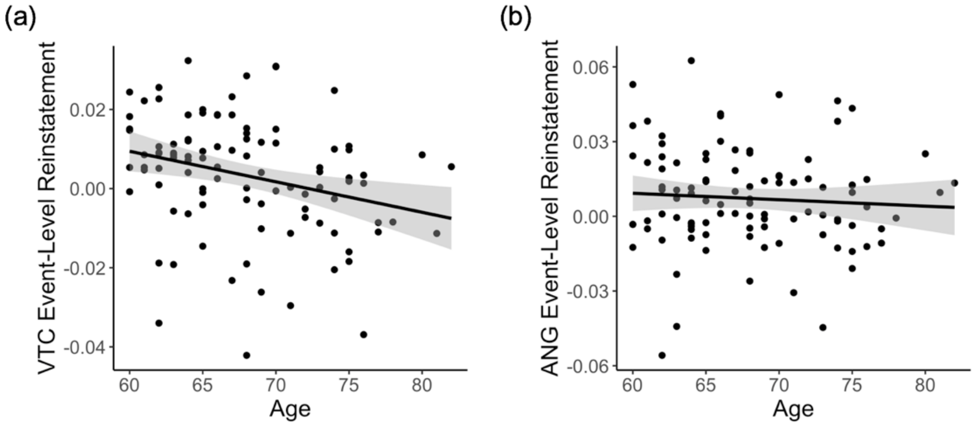


*Figure S6. Event-Level Reinstatement Strength (within-event – within-category ERS) Declines with Age in (a) VTC , but not (b) ANG during Associative Hits. Each point on the scatterplot represents an individual subject. Plots also show the linear model predictions (black line) and 95% confidence interval (shaded area). VTC = ventral temporal cortex; ANG = angular gyrus.*

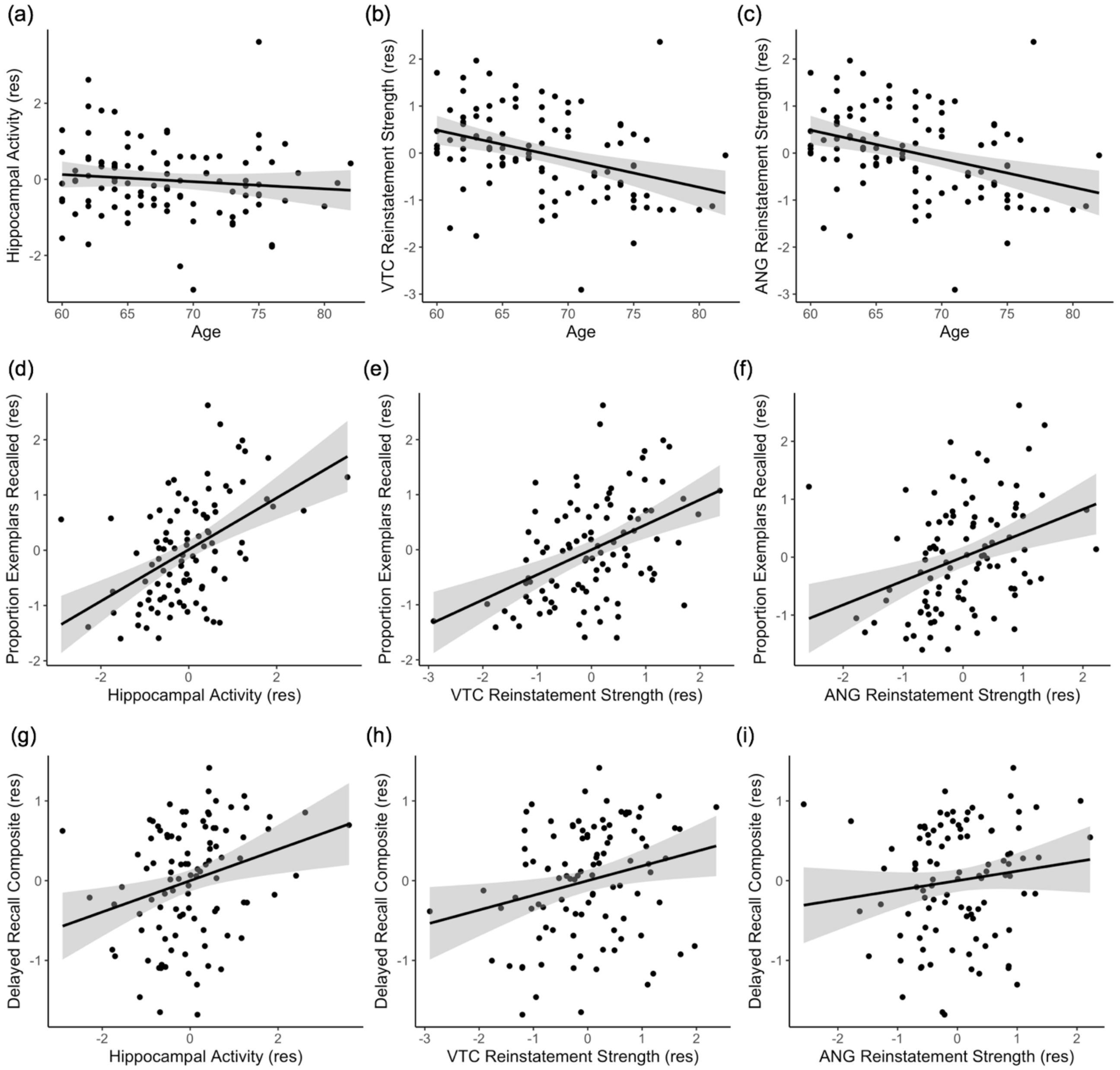


*Figure S7. Partial Plots of Individual Differences in Pattern Completion Assays. (a-c) Effects of age on hippocampal activity (associative hits – correct rejections) and reinstatement strength (mean logits) in VTC and ANG during associative hits. (d-f) Independent of age, individual differences in hippocampal activity and reinstatement strength in VTC and ANG during associative hits significantly predict exemplar-specific recall. (g-i) Independent of age, individual differences in hippocampal activity and VTC reinstatement strength also explain significant variability in standardized delayed recall performance; the relation with ANG reinstatement did not reach significance. Scatterplots reflect partial plots controlling for relevant nuisance variables. Each point represents an individual participant. Plots also show linear model predictions (black line) and 95% confidence intervals (shaded area). VTC = ventral temporal cortex; ANG = angular gyrus.*

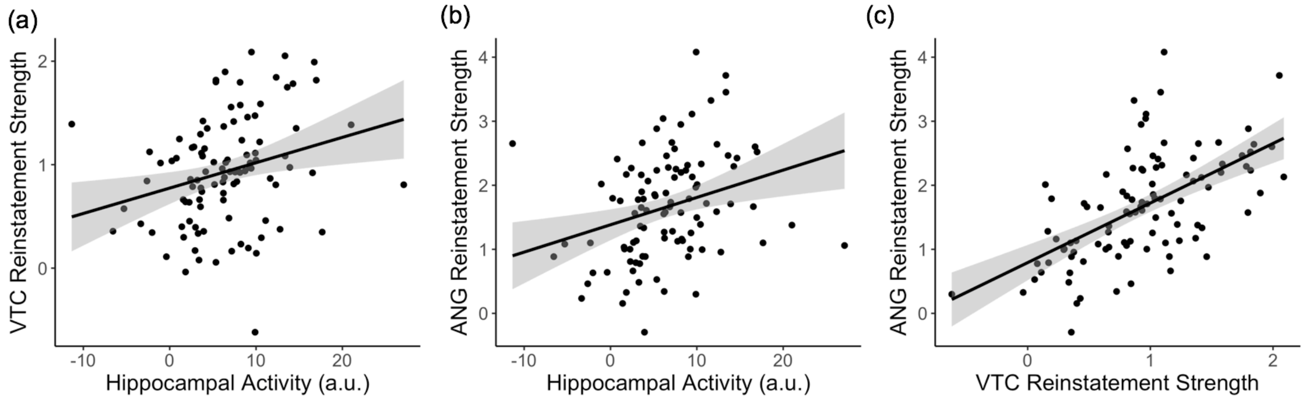


*Figure S8. Pattern Completion Metrics Covary across Participants. Hippocampal activity during associative hits (associative hits – correct rejections) predicts category-level reinstatement strength (logits) in (a) VTC and (b) ANG during associative hits. (c) Category-level reinstatement strength (logits) in VTC and ANG during associative hits are related. Each point on the scatterplot represents an individual subject. Plots also show the linear model predictions (black line) and 95% confidence interval (shaded area). VTC = ventral temporal cortex; ANG = angular gyrus.*

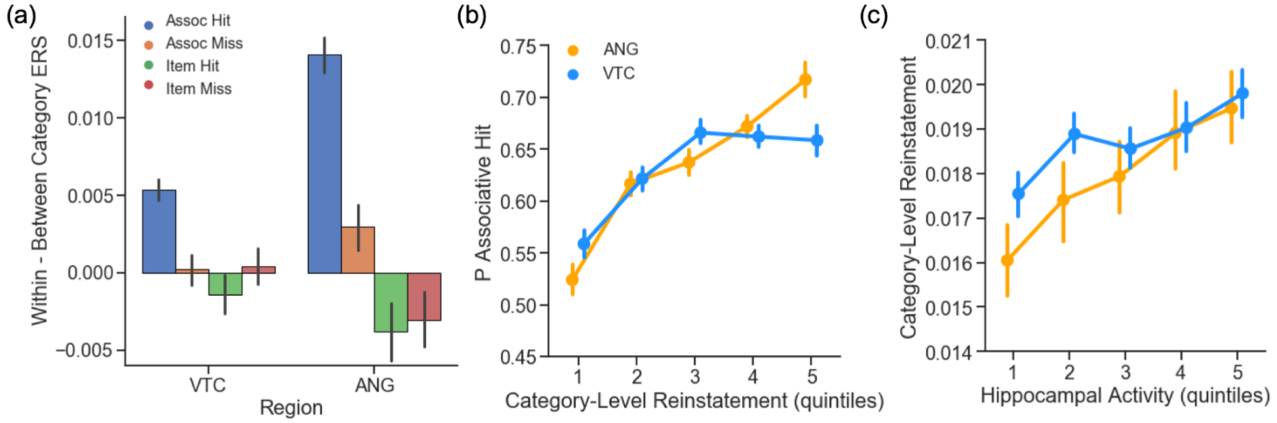


*Figure S9. Category Reinstatement Effects Computed via Encoding-Retrieval Similarity. (a) Within-category ERS (Pearson r) was greater than between-category ERS during successful, but not unsuccessful, associative retrieval. (b) Trial-wise within-category reinstatement strength in VTC and ANG predicted the probability of an associative hit. (c) Trial-wise increases in hippocampal activity predicted greater within-category reinstatement strength in VTC and ANG. For visualization, data for each participant were binned into quintiles based on (b) ERS and (c) hippocampal activity. Statistics were conducted on trial-wise data, z-scored within participant. Error bars represent standard error of the mean. ERS = Encoding Retrieval Similarity. VTC = ventral temporal cortex; ANG = angular gyrus.*

**Supplementary Results**

*Behavioral Data*

On average, associative *d'* was better for words paired with faces (M = 2.16, SD = .68) than words paired with places (M = 1.85, SD = .74; $t(99) = 5.37$, $p < 10^{-7}$). Median RTs varied as a function of response type, such that associative hits (M = 2382ms, SD = 421ms) were faster than associative misses (M = 2939ms, SD = 553ms; $t(99)=12.09$, $p < 10^{-16}$), item only hits (M = 3198ms, SD = 600ms; $t(99)=13.80$, $p < 10^{-16}$), and item misses (M = 2689ms, SD = 568ms ; $t(99)=4.99$, $p < 10^{-6}$). Associative hit RT was significantly faster for face trials (M = 2271ms, SD = 497ms) than place trials (M = 2548ms, SD = 413ms; $t(99)=8.40$, $p < 10^{-13}$).

*Effects of Motion*

To investigate possible effects of head motion on the primary variables of interest in our between-subject analyses, we computed mean framewise displace, separately for study and test runs. There was a trend towards an age-related increase in head motion (study: $\beta = .19$, $p = .05$; test: $\beta = .15$, $p = .14$), but no significant effect of sex (study: $\beta = -.30$, $p = .15$; test: $\beta = .29$, $p = .16$), and no effect of education (all *p*s > .61). While head motion was related to encoding classifier performance (VTC: $\beta = -.48$, $p < 10^{-7}$; ANG: $\beta = -.34$, $p < .001$), no significant relationship emerged between head motion and (a) category reinstatement strength (VTC: $\beta = -.06$, $p > .52$; ANG: $\beta = -.19$, $p > .06$), (b) event-level reinstatement strength (VTC: $\beta = .02$, $p = .34$; ANG: $\beta = .03$, $p = .13$), nor (c) hippocampal activity ($\beta = -.16$, $p > .10$). Head motion was unrelated to associative *d'* nor exemplar-specific recall (all *p*s > .55). To control for possible effects of head motion on our variables of interest, we adjusted each variable to reflect residuals from the regression models described here.

*Effects of Encoding Classifier Strength on Reinstatement Strength*

In our primary analyses (**Main Text**), we control for encoding classifier strength in all models in which reinstatement strength a) predicted behavioural variables (memory accuracy, RT), and b) was the dependent variable, to account for possible effects of encoding strength on subsequent reinstatement strength. At the trial-level, linear mixed effects models revealed that encoding classifier strength of individual items predicted later reinstatement strength (VTC: $\chi^2(1) = 13.96$, $p < .001$; ANG: $\chi^2(1) = 30.16$, $p < 10^{-8}$). Individual differences analyses similarly revealed that encoding strength (mean logits across leave-one-run-out-n-fold cross validation) was a significant predictor of reinstatement strength during associative hits (mean logits; VTC: $\beta = 0.45$, $p < 10^{-5}$ ; ANG: $\beta = 0.62$, $p < 10^{-11}$; **Figure S1**). In all multiple regression analyses, reinstatement strength is adjusted by encoding strength using residuals from these models.

*Trial-wise Analyses*

*Pattern Similarity Analysis: Category-Level Reinstatement*

To complement the classification results, we also examined category-level cortical reinstatement using an encoding-retrieval similarity (ERS) analysis approach. Evidence for category-level reinstatement was considered present when within-category ERS was significantly greater than between-category ERS. Category-level reinstatement was robust during associative hits in VTC ($t(99) = 8.84$, $p < 10^{-14}$) and ANG ($t(99) = 12.15$, $p < 10^{-16}$), but was not observed during associative misses (VTC: $p = .97$; ANG: $p = .19$), item only hits (VTC: $t(51) = -1.22$, $p = .23$; ANG: $t(51) = -1.92$, $p = .06$), nor item misses (VTC: $p = .69$; ANG: $t(83) = -1.89$. $p = .06$; Figure S9a). Category-level reinstatement was stronger in ANG than VTC ($t(99) = 9.0$, $p < 10^{-14}$).

We next assessed whether trial-wise within-category ERS predicts the probability of an associative hit using logistic mixed effects models, with random intercepts for subjects and random slopes for the effects of interest, separately for each ROI. Greater within-category ERS predicted an increased probability of an associative hit (VTC: $\chi^2(1) = 18.91$, $p < 10^{-5}$; ANG: $\chi^2(1) = 53.05$, $p < 10^{-13}$; Figure S9b). In VTC, this relationship varied marginally by stimulus category ($\chi^2(1) = 4.14$, $p < .05$), being stronger for place trials ($\chi^2(1) = 24.03$, $p < 10^{-7}$) than face trials ($\chi^2(1) = 5.48$, $p < .05$). In ANG, this relationship did not vary by stimulus category ($p$ > .581). A relationship between within-category ERS and RT during associative hits was not significant in VTC ($p$ = 0.110) or ANG ($p$ = .946).

Finally, when considering all trials, hippocampal activity predicted the magnitude of within-category ERS in both regions (VTC: $\chi^2(1) = 8.65$, $p = .003$; ANG: $\chi^2(1) = 9.22$, $p = .002$; Figure S9c). This relationship did not differ by category (VTC: $p$ = .905; ANG: $p$ = .538). However, when considering associative hit trials only, hippocampal activity was no longer a significant predictor of within-category ERS in VTC ($p$ = .162) or ANG ($p$ = .362), nor did this effect interact with stimulus category (VTC: $p$ = .657; ANG: $p$ = .841).

*Hippocampal Subfield Activity at Retrieval Predicts Behaviour and Cortical Reinstatement*

Having observed that hippocampus activity predicts associative retrieval and the strength of cortical reinstatement (see **Main Text**), we assessed whether this effect varied across hippocampal subfields (Figure S5). Within the body of the hippocampus, activity in each subfield significantly predicted associative retrieval (DG/CA3: $\chi^2(1) = 37.03$, $p < 10^{-9}$; CA1: $\chi^2(1) = 23.16$, $p < 10^{-6}$; Sub: $\chi^2(1) = 15.29$, $p < 10^{-5}$). We additionally observed an interaction between univariate activity and subfield ($\chi^2(1) = 12.99$, $p < .001$), such that DG/CA3 activity was a significantly stronger predictor than CA1 ($z = 2.31$, $p < .05$) or Sub ($z = 3.55$, $p < .001$), whereas CA1 and Sub did not significantly differ ($z = 1.24$, $p > .21$) (Figure S5a). Additionally, activity in all three subfields predicted VTC category reinstatement strength when considering all trials (DG/CA3: $\chi^2(1) = 12.25$, $p < .001$; CA1: $\chi^2(1) = 15.40$, $p < 10^{-5}$; Sub: $\chi^2(1) = 13.38$, $p < .001$; here the interaction was not significant, all $p$ = .703; Figure S5b). Subfield activity no longer significantly predicted VTC reinstatement when considering associative hit trials only (DG/CA3: $p$ = .748; CA1: $p$ = .175; Sub: $p$ = .504). Similarly, category reinstatement strength in ANG was predicted by DG/CA3 ($\chi^2(1) = 7.89$, $p = .005$) and CA1 activity ($\chi^2(1) = 10.49$, $p = .001$), but not Sub ($p$ = .458) activity; the DG/CA3 and CA1 effects did not differ ($p$ = .988; Figure S5c). When considering associative hit trials, only DG/CA3 remained marginally significant ($\chi^2(1) = 2.86$, $p$ =.091; CA1: $p$ = .437; Sub: $p$ = .572).

We additionally examined relationships between univariate activity and (a) memory and (b) reinstatement strength in the head and tail of the hippocampus, two areas where subfields could not be delineated at the present image resolution (76). We observed an interaction by subregion ($\chi^2(1) = 31.44$, $p < 10^{-8}$), such that this relationship was stronger in the hippocampal head: $\chi^2(1) = 65.76$, $p < 10^{-16}$ ) than hippocampal tail ($\chi^2(1) = 24.08$, $p < 10^{-7}$). Similarly for reinstatement strength, we observed an interaction by subregion (VTC: $\chi^2(1) = 14.86$, $p < .001$; ANG: $\chi^2(1) = 24.77$, $p < 10^{-8}$), reflecting a stronger relationship between univariate activity and reinstatement strength in the hippocampal head (VTC: $\chi^2(1) = 44.97$, $p < 10^{-11}$; ANG: $\chi^2(1) = 46.76$, $p < 10^{-12}$) than tail (VTC: $\chi^2(1) = 44.97$, $p = .021$; ANG: $p$ =.743). When considering associative hit trials only, only activity in the hippocampal head remained significant (VTC: $\chi^2(1) = 11.70$, $p < .001$; ANG: $\chi^2(1) = 20.35$, $p < 10^{-6}$; tail: VTC: $p = .762$; ANG: $p > .358$). Together, these results suggest a greater contribution of the hippocampal head than the hippocampal tail to associative retrieval and to driving cortical reinstatement in the present paradigm.

*Reinstatement Strength vs Memory Behaviour: Effect of Stimulus Category*

To assess possible effects of stimulus category on the relationship between category-level reinstatement strength and the probability of associative retrieval success, we re-ran models including an interaction term between category and reinstatement strength. The relationship between category-level reinstatement and the probability of associative retrieval varied by category (VTC: $\chi^2(1) = 29.55$, $p < 10^{-8}$; ANG: $\chi^2(1) = 9.63$, $p = .002$). The relationship was present for both face and place trials across regions, but was stronger in VTC on place trials ($\chi^2(1) = 75.81$, $p < 10^{-18}$) than face trials ($\chi^2(1) = 21.89$, $p < 10^{-6}$) and stronger in ANG on face trials ($\chi^2(1) = 93.73$, $p < 10^{-22}$) than place trials ($\chi^2(1) = 55.84$, $p < 10^{-14}$). With respect to decision RT, we observed an interaction between stimulus category and classifier evidence in both VTC ($\chi^2(1) = 9.39$, $p = .002$) and ANG ($\chi^2(1) = 92.29$, $p < 10^{-22}$). In VTC, this effect was stronger on face trials ($\chi^2(1) = 36.40$, $p < 10^{-9}$) than place trials ($\chi^2(1) = 8.46$, $p = .004$). In ANG, greater reinstatement was associated with faster responses on face trials ($\chi^2(1)$ =

45.51, $p < 10^{-11}$), but the opposite was true on place trials ($\chi^2(1) = 10.43$, $p = .001$). The relationship between event-level reinstatement and the probability of an associative hit did not significantly vary according to stimulus category (VTC: $\chi^2(1) = 2.17$, $p = .141$; ANG: $p = .763$).

*Hippocampal Activity, Reinstatement, & Memory Behaviour: Effect of Stimulus Category*

The relationship between hippocampal activity and associative retrieval success did not significantly vary with the category of the retrieved associate ($\chi^2(1)= 2.86$, $p = .091$). Similarly, the relationship between hippocampal activity and associative hit decision RT did not vary with category ($p = .905$). The relationship between hippocampal activity and category-level reinstatement strength was modulated by stimulus category (VTC: $\chi^2(1)= 235.87$, $p < 10^{-42}$ ; ANG: $\chi^2(1)= 8.47$, $p = .004$). In ANG, this relationship was stronger on face trials ($\chi^2(1)= 28.28$, $p < 10^{-7}$) than place trials ($\chi^2(1)= 3.15$, $p = .077$), whereas in VTC this relationship was greater on place trials ($\chi^2(1)= 82.93$, $p < 10^{-20}$) than face trials ($\chi^2(1)= 11.96$, $p < .001$). The relationship between hippocampal activity and event-level reinstatement strength in VTC was also modulated by stimulus category ($\chi^2(1)= 4.62$, $p = .031$), such that this effect was significant on place trials ($\chi^2(1)= 9.13$, $p = .003$), but not on face trials ($p = .88$). There was no interaction between hippocampal activity and event-level reinstatement strength in ANG ($\chi^2(1)= 2.23$, $p = .135$).

*Individual Difference Analyses*

*Relationship between Subject-level Indices of Pattern Completion*

We considered the relationships between our measures, controlling for the effect of age. We found that: a) hippocampal activity during associative hits (associative hit – CR) significantly predicted category-level reinstatement (i.e., mean logits) during associative hits in both VTC ($\beta = .19$, $p < .05$; Figure S7a) and ANG ($\beta = .21$, $p < .01$; Figure S7b); b) the relationship between hippocampal activity and event-level reinstatement (i.e., within-event – between category ERS) during associative hits was not significant in either region (*p*s > .21); and c) category-level reinstatement and event-level reinstatement in VTC predicted that in ANG (category-level: $\beta = .44$, $p < 10^{-7}$; Figure S7c; ERS: $\beta = .54$, $p < 10^{-9}$). Taken together, these results provide evidence that, in older adults, hippocampal retrieval activity and cortical reinstatement — putative assays of pattern completion — co-vary with one another, even after controlling for the effects of age.

*Neural Predictors of Individual Differences in Associative d'*

To complement our analyses relating subject-level hippocampal and cortical estimates of pattern completion to individual differences in post-scan exemplar-specific recall (see **Main Text**), we conducted the same analyses with respect to associative *d'*. Category reinstatement strength in VTC partially mediated the relationship between age and associative *d'* (total effect = -0.32, z = -3.42, $p < 0.001$; direct effect = -0.18, z = -1.94, $p = .05$; indirect effect = -0.14, z = -2.86, $p < .005$, 95% CI = -0.241, -0.045). This effect was marginally significant in ANG (total = -0.32, z = -3.42, $p < 0.001$; direct = -0.26, z = -2.88, $p < .005$; indirect = -0.06, z = -1.71, $p = .09$, 95% CI = -0.131, 0.009).

Independent of age, separate regression models for each pattern completion metric (adjusted by relevant nuisance variables) revealed that individual differences in associative *d'* were predicted by hippocampal activity ($\beta = 0.34$, $p < .001$) and category-level reinstatement strength during associative hits (VTC: $\beta = 0.38$, $p < .001$; ANG: $\beta = 0.39$, $p < .001$). In contrast, individual differences in event-level reinstatement did not explain significant variance in associative *d'* (all *p*s > .14). Hierarchical regression further revealed that age, hippocampal activity, and reinstatement strength in VTC explained unique variance in associative *d',* whereas the inclusion of ANG reinstatement did not explain additional variance (see **Table S1**). Thus, individual differences in hippocampal univariate activity and cortical reinstatement strength measures provide complementary information in explaining individual differences in associative memory, whereas indices of reinstatement strength explain shared variance.

*Table S1: Summary of Regression Analysis Predicting Associative d'*

| | Variable | β | SE | p | Adjusted $R^2$ |
|---|---|---|---|---|---|
| Step 1 | Age | -0.324 | 0.096 | 0.001*** | 0.096 |
| Step 2 | Age | -0.283 | 0.090 | 0.001*** | 0.205 |
| | Hippocampal $\text{Activity}_{adj}$ | 0.349 | 0.091 | 0.001*** | |
| Step 3a | Age | -0.169 | 0.090 | 0.063~ | 0.295 |
| | Hippocampal $\text{Activity}_{adj}$ | 0.284 | 0.088 | 0.002** | |
| | VTC $\text{Reinstatement}_{adj}$ | 0.368 | 0.100 | 0.001*** | |
| Step 3b | Age | -0.245 | 0.089 | 0.007** | 0.249 |
| | Hippocampal $\text{Activity}_{adj}$ | 0.282 | 0.093 | 0.003** | |
| | ANG $\text{Reinstatement}_{adj}$ | 0.303 | 0.117 | 0.011* | |
| Step 4 | Age | -0.179 | 0.090 | 0.064~ | 0.297 |
| | Hippocampal $\text{Activity}_{adj}$ | 0.262 | 0.089 | 0.004*** | |
| | VTC $\text{Reinstatement}_{adj}$ | 0.310 | 0.113 | 0.007*** | |
| | ANG $\text{Reinstatement}_{adj}$ | 0.146 | 0.127 | 0.254 | |
| Step 5 | Age | -0.119 | 0.090 | 0.190 | 0.336 |
| | Hippocampal $\text{Activity}_{adj}$ | 0.228 | 0.088 | 0.012* | |
| | VTC $\text{Reinstatement}_{adj}$ | 0.312 | 0.100 | 0.002** | |
| | Delayed Recall | 0.328 | 0.124 | 0.009** | |

*Note*. Adj = Adjusted by relevant nuisance regressors; SE= standard error; VTC = ventral temporal cortex; ANG = angular gyrus; ~ p < 0.1, * p < 0.05, ** p < .01, *** p < .001 **** p < $10^{-5}$

*Hippocampal Subfield Activity Predicts Individual Differences in Associative Memory*

Finally, we assessed whether the relationship between hippocampal activity during associative retrieval (associative hit – CR) and memory was specific to a particular subfield of the hippocampus. Separate regression models for each subfield, each controlling for age and head motion revealed that individual differences in activity in all three subfields significantly predicted associative *d'* (DG/CA3: $\beta$ = .26, *p* < .01; CA1: $\beta$ = .20, *p* < .05; Sub: $\beta$ = .19, *p* < .05) and exemplar-specific recall (DG/CA3: $\beta$ = .32, *p* < .005; CA1: $\beta$ = .28, *p* < .005; Sub: $\beta$ = .28, *p* < .01). When all body subfields were considered in the same model, only DG/CA3 remained marginally significant (associative *d'*: DG/CA3: $\beta$ = .19, *p* = .07; CA1: $\beta$ = .09, *p* = .40; Sub: $\beta$ = .08, *p* = .45; exemplar-specific recall: DG/CA3: $\beta$ = .21, *p* < .05; CA1: $\beta$ = .15, *p* = .14; Sub: $\beta$ = .16, *p* = .13), but the magnitude of this effect did not differ across subfields (all $p > .52$). Moreover, activity in the hippocampal head and hippocampal tail both significantly predicted associative *d'* (head: $\beta = .288$, $p < .005$; tail: $\beta = .256$, $p < .01$) and exemplar-specific recall (head: $\beta = .391$, $p < 10^{-5}$; tail: $\beta = .246$, $p < .01$). Activity in both regions remained significant when included in the same model (associative *d'*: head: $\beta = .237$, $p < .05$; tail: $\beta = .192$, $p < .05$; exemplar-specific recall: head: $\beta = .350$, $p < .001$; tail: $\beta = .153$, $p = .08$). These results suggest that the positive relationship between hippocampal activity during memory retrieval and associative memory is qualitatively similar across hippocampal subfields in putatively healthy older adults.
